# A cell atlas of the larval *Aedes aegypti* ventral nerve cord

**DOI:** 10.1101/2023.09.08.556941

**Authors:** Chang Yin, Takeshi Morita, Jay Z. Parrish

**Affiliations:** Department of Biology, University of Washington, Seattle, WA 98195, USA; Division of Education, Marine Biological Laboratory, 7 MBL Street, Woods Hole, MA 02543, USA; Laboratory of Neurogenetics and Behavior, The Rockefeller University, New York, NY 10065, USA; Howard Hughes Medical Institute, New York, NY 10065, USA

## Abstract

Mosquito-borne diseases account for nearly one million human deaths annually, yet we have a limited understanding of developmental events that influence host-seeking behavior and pathogen transmission in mosquitoes. Mosquito-borne pathogens are transmitted during blood meals, hence adult mosquito behavior and physiology have been intensely studied. However, events during larval development shape adult traits, larvae respond to many of the same sensory cues as adults, and larvae are susceptible to infection by many of the same disease-causing agents as adults. Hence, a better understanding of larval physiology will directly inform our understanding of physiological processes in adults. Here, we use single cell RNA sequencing (scRNA-seq) to provide a comprehensive view of cellular composition in the *Aedes aegypti* larval ventral nerve cord (VNC), a central hub of sensory inputs and motor outputs which additionally controls multiple aspects of larval physiology. We identify more than 35 VNC cell types defined in part by neurotransmitter and neuropeptide expression. We also explore diversity among monoaminergic and peptidergic neurons that likely control key elements of larval physiology and developmental timing, and identify neuroblasts and immature neurons, providing a view of neuronal differentiation in the VNC. Finally, we find that larval cell composition, number, and position are preserved in the adult abdominal VNC, suggesting studies of larval VNC form and function will likely directly inform our understanding adult mosquito physiology. Altogether, these studies provide a framework for targeted analysis of VNC development and neuronal function in *Aedes* larvae.

## Introduction

Mosquito-borne diseases afflict nearly 700 million people annually resulting in more than 700,000 deaths, making mosquitoes the deadliest animal in the world (WHO 2020). This disease burden is primarily attributable to two mosquito species: *Anopholes gambii*, which transmit malaria in tropical climates, and *Aedes aegypti* which inhabit cooler, urbanized environments where they transmit chikungunya, dengue, yellow fever, zika, and other viruses. Urbanization and climate change are intensifying risks of human exposure to these viruses by expanding mosquito habitats, accelerating mosquito development, and influencing pathogen reproduction rates (Liu-Helmersson et al. 2014; Kraemer et al. 2019). However, despite the enormous impact of mosquitos on human health, we have a limited understanding of developmental events that influence host-seeking behavior and pathogen transmission in mosquitos.

Like other holometabolous insects, mosquito development includes distinct juvenile forms – embryo, four larval instars and pupa – and aspects of juvenile development directly impact adult traits. First, larvae exhibit many of the same behaviors, respond to many of the same chemical/sensory cues, and are themselves susceptible to infection by many of the same disease-causing agents as adults (Bara et al. 2013). Hence, a better understanding of larval physiology will directly inform our understanding of physiological processes in adults. Second, many larval structures likely persist in adults. Whereas many dipterans undergo protracted metamorphosis during which their adult nervous system is produced through extensive remodeling and replacement of larval neurons, *Aedes aegypti* pupae complete metamorphosis in <36 h and *Anopholes gambii* can complete pupal development in under a day (Batume et al. 2022). This rapid metamorphosis likely constrains the extent of nervous system remodeling. Indeed, the limited number of longitudinal studies reported to date are consistent with the model that many larval neurons persist into adult mosquitos. For example, larvae exhibit well-developed optic lobes that resemble adult structures, and identified serotonergic neurons persist into adults (Moffett and Moffett 2005; Mysore et al. 2011). Third, events in larval development including diet, thermal cues, intraspecies competition, and other environmental conditions directly impact pathogen infection and vectorial capacity of adults (Alto et al. 2008; Lambrechts et al. 2011; Moller-Jacobs et al. 2014; Dickson et al. 2017; Herd et al. 2021). Finally, targeting larvae for population control provides potential advantages including the organism’s limited range and opportunities to intervene prior to reproductive maturity.

*Aedes aegypti* (subsequently referred to as *Aedes*) mosquitos have a segmented central nervous system comprised of the supraesophageal ganglion (brain) and subesophageal ganglion located in the head, and the ventral nerve cord (VNC) spanning the thorax and abdomen. Portions of the adult mosquito nervous system have been extensively studied (Matthews et al. 2016; Cui et al. 2022; Herre et al. 2022), but comprehensive studies linking gene expression, morphology, and physiology are largely lacking for larvae. Inspired by the utility of scRNA-seq cell atlases in other organisms (Davie et al. 2018; Tasic et al. 2018; Brunet Avalos et al. 2019; Berg et al. 2021) we set out to generate a cell atlas of the *Aedes* larval nervous system, focusing on the VNC which serves as a nexus for controlling mosquito behavior, physiology and developmental progression. The VNC receives sensory inputs which it integrates with descending inputs from the brain, contains the motor neurons responsible for locomotor actions, contains monoaminergic neurons that shape behavioral states, and additionally contains peptidergic neurons that control developmental timing, metabolism, ion homeostasis, circadian rhythms, and behavior (Santos et al. 2007). In addition, the larval VNC has been extensively characterized in other insects, most notably *Drosophila melanogaster*, providing a framework for defining the developmental origin, transcriptional profile, and functional properties of *Aedes* counterparts.

In this study, we provide a detailed view of the cellular composition of the *Aedes* larval VNC. First, we anatomically define the cellular repertoire of the VNC. This analysis revealed that the *Aedes* larval VNC is comprised of roughly half as many cells as its *Drosophila* counterpart. Second, we generated an atlas of VNC neurons containing 6564 single-cell transcriptional profiles that we utilized to identify more than 35 cell types. We find that patterns of neurotransmitter and neuropeptide expression are defining features of many of these cell types, and although the individual fast-acting neurotransmitters (FANs) acetylcholine, GABA, and glutamate are expressed in roughly equivalent numbers of cells, we find that GABAergic neurons are concentrated in a posterior domain of each ganglion. To illustrate the utility of our dataset in identifying features of cell types, we transcriptionally and anatomically define different classes of monoaminergic and peptidergic neurons and identify potential regulators of their diversity. We additionally capture transcriptional profiles of putative neuroblasts and immature neurons, providing insight into transcriptional mechanisms of neuronal differentiation and post-embryonic VNC development in *Aedes*. Finally, we explore the relationship between the larval and adult VNC and find that larval neurons largely persist in the same number and position, further underscoring the significance of understanding larval VNC development and function.

## Results

### Single-cell transcriptomic atlas of the larval ventral ganglia

The *Aedes* VNC consists of two morphologically distinct segmental ganglia: a hemi-fused ganglion containing the three thoracic neuromeres and a chain of eight uniform segmental units distributed throughout the abdomen (Fig. 1A). Although the thoracic units are larger than those in abdominal segments, each ganglion exhibits the same basic structure: a bilaterally symmetric cortex of cell bodies surrounding a central zone of neuropile (Fig. 1B). As a first approach to studying cellular complexity in the larval VNC we monitored the cellular composition of each neuromere using the DNA dye Hoechst to label nuclei of all cells and antibodies to Elav, a pan-neuronal marker in *Drosophila* and mosquitos (Robinow and White 1991; Raji and Potter 2021) that labels both immature and mature neurons (Koushika et al. 1996). For these studies we chose two timepoints spanning the developmental window during which post-embryonic neurogenesis occurs in *Drosophila*: day 3 (L2 stage) and day 5 (L4 stage) of larval development (Truman and Bate 1988). Our analysis revealed several features of VNC organization. First, thoracic ganglia had more cells on average than abdominal ganglia, consistent with the larger overall size of the thoracic ganglia (Fig. 1C). Second, at each timepoint the average number of Hoechst-positive nuclei and Elav-positive neurons was comparable in each thoracic neuromere; the same was true in abdominal neuromeres (Fig. 1C, 1D). In sum, the L4 *Aedes* larval VNC is comprised of ∼3500 cells, including ∼2500 neurons. Third, neither thoracic nor abdominal ganglia exhibited any significant increase in the number of neurons or cells overall during this time window. Instead, thoracic ganglion exhibited a significant reduction in overall cell number which predominantly reflected a reduction in the number of Elav-positive neurons. Finally, although Elav-positive cells were the principal cell type in all segments, thoracic ganglia exhibited a smaller proportion of Elav-positive cells (63% in T1, 78% in A1; Fig. 1E), indicating that they contained a larger proportion of glia and/or neuroblasts. To directly evaluate glial composition of each ganglion we stained larval VNCs with antibodies to *Drosophila* glial markers including the pan-glial marker repo and markers for wrapping glia and astrocytes, but we were unable to identify antibodies that yielded specific signal (Table S1).

**Figure 1.**
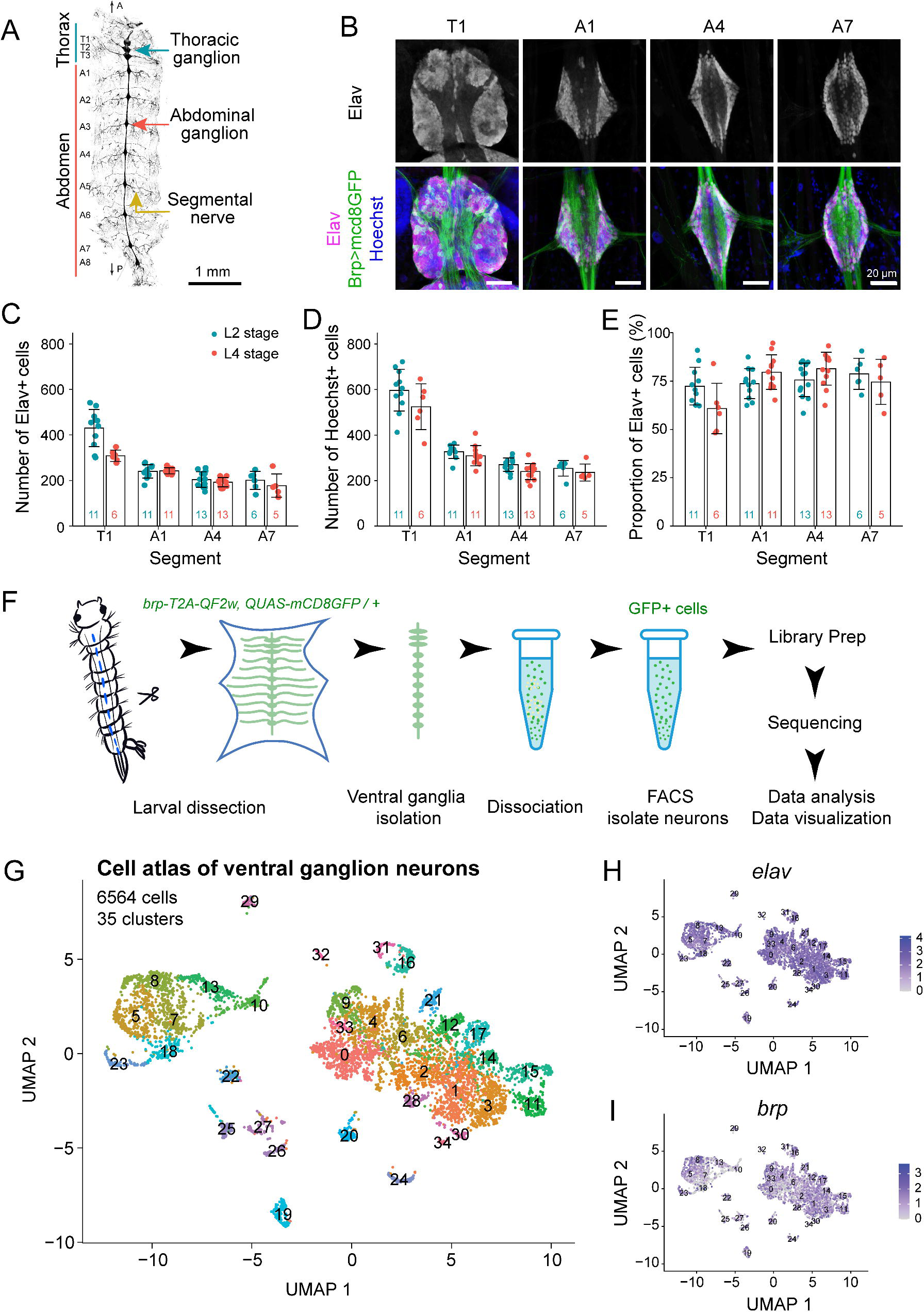
Morphology, cellular composition, and single-cell RNA-seq atlas of *Aedes* larval VNC. (A) Structure of larval VNC and segmental nerves visualized in a fillet preparation of a *brp-T2A-QF2w / +; QUAS-mcd8GFP / +* larva. (B) Cellular composition of thoracic and abdominal ganglia. Images show maximum projections of representative confocal stacks taken from segments T1, A1, A4 and A7 of larvae stained with Hoechst to label all nuclei together with antibodies to Elav to label neuronal nuclei and GFP to label neuronal membranes. Images depict Elav+ cells (top) and a montage of all three markers (bottom). (C-E) Quantification of ganglion cell numbers. Plots depict the number of (C) Elav-positive cells, (D) Hoechst-positive cells, and (E) the proportion of cells which were Elav-positive in four representative segments (T1, A1, A4, A7) at two different larval time points (3 days after hatching and 5 days after hatching). Bars depict mean values, points represent measurements from individual ganglia, sample numbers are indicated for each segment/time point combination, and error bars depict standard deviation. *P<0.05, two-way ANOVA followed by Fisher’s LSD test. The two-way ANOVA revealed a statistically significant interaction between the effects of developmental time and segment position on neuron number (F(3, 69) = 9.846, p < 0.0001) and the proportion of neurons (F(3, 68) = 3.756, p = 0.0148). Simple main effects analysis showed that developmental time had statistically significant effects on cell number (p < 0.0001), neuron number (p < 0.0001), and neuron proportion (p = 0.0012), and that segment position had statistically significant effects on cell number (p = 0.0112) and neuron number (p = 0.0003). (F) Schematic workflow for generating single cell transcriptional profiles. Mosquito larval VNCs were isolated and dissociated prior to sorting GFP-positive neurons by FACS. Following library preparation, sequencing, and quality-based filtering, clustering analysis was used to identify gene expression relationships between profiled cells. (G) *Aedes* larval VNC cell atlas. The UMAP plot depicts 6,564 neurons grouped into 35 clusters. (H-I) Feature plots depicting neuronal marker gene expression across the *Aedes* VNC cell atlas. Genotype for all panels: *brp-T2A-QF2w / +; QUAS-mcd8GFP / +*.

To define the neuronal repertoire of the larval VNC, we dissociated larval VNCs expressing *QUAS-mCD8-GFP* under the control of the pan-neuronal QF driver, *brp-T2A-QF2w* (Zhao et al. 2021) from fourth instar larvae and subjected 10,000 FACS-sorted single neurons to single-cell RNA sequencing using droplet microfluidics (10x Chromium) (Fig. 1F). After stringent quality-based filtering according to the number of unique molecular identifiers (UMIs), the number of genes, the proportion of mitochondrial genes, and the proportion of ribosomal genes (Fig. S1) we used graph-based methods to cluster the remaining 7965 cells based on shared gene expression programs (Hao et al. 2021). This initial clustering analysis yielded 25 cell clusters (Fig. S2) which were evaluated according to several criteria including cell number, cell quality, and marker gene expression. Among these clusters, we identified two heterogeneous clusters with low overall cell counts; these clusters were eliminated from further analysis. We additionally identified a cluster that broadly expressed the pan-glial marker gene *repo* and contained subsets of cells that expressed markers for glial subtypes including wrapping glia, perineural glia, and astrocytes (Fig. S2); we identified marker genes for this cluster to facilitate future studies of larval glia (Table S2). However, this cluster had lower numbers of genes and total counts per cell compared to the neuronal clusters and additionally contained cells that expressed neuronal marker genes (*Syt*, *elav*, and *brp*), hence we eliminated the cluster from further analysis.

Reclustering analysis of the remaining 6564 cells identified 35 discrete clusters that populate the VNC (Fig. 1G) and express a battery of known pan-neuronal marker genes including *embryonic lethal abnormal vision* (*elav*), *bruchpilot* (*brp*), *Synaptotagmin* (*Syt*), *Cadherin-N* (*CadN*), and *neuronal Synaptobrevin* (*nSyb*) (Fig 1H-1I; Fig. S3). A recent study of cellular diversity in the adult *Aedes* brain identified comparable levels of cellular diversity using single nuclei (sNuc)-Seq (Cui et al. 2022); however, the shallow depth (1-10k reads/cell, compared to 100k reads/cell in our atlas) and limited coverage offered by sNuc-Seq precludes meaningful comparison of the identified subsets.

### Neurotransmitter expression as a marker for larval VNC cluster identity

The repertoire of neurotransmitter(s) expressed by a neuron is a diagnostic feature of neuronal type, therefore as a first measure of functional diversity in the *Aedes* VNC we examined marker gene expression for classical neurotransmitters, including fast acting neurotransmitters (FANs) and monamines. We assigned neurotransmitter identity to cells based on expression of the following genes involved in transmitter biosynthesis or vesicular loading: *vesicular glutamate transporter* (*VGlut*) and *Glutamic acid decarboxylase 1* (*Gad1*) for glutamatergic (Glu) neurons; *Vesicular GABA transporter* (*GAT*) for GABAergic neurons; *Vesicular acetylcholine transporter* (*VAChT*) and *Choline acetyltransferase* (*ChAT*) for cholinergic (ACh) neurons; *vesicular monoamine transporter* (*Vmat*) as a marker for all monoaminergic neurons; *Tyramine beta hydroxylase* (*Tbh*) for octopaminergic neurons; *serotonin transporter* (*SerT*) for serotonergic neurons; *tyrosine hydroxylase* (*TH*) and *Dopamine transporter* (*Dat*) for dopaminergic neurons.

Consistent with similar studies in *Drosophila* (Vaaga et al. 2014; Croset et al. 2018; Brunet Avalos et al. 2019; Allen et al. 2020; Corrales et al. 2022), we found that *Aedes* larval VNC clusters exhibited largely non-overlapping expression of FAN markers: in 28/35 clusters, >70% of cells within that cluster expressed marker genes for a particular FAN (Fig. 2A, S4, Table S3), corresponding to 13 clusters of ACh neurons, 7 clusters of GABA neurons, and 8 clusters of Glu neurons. Although these clusters expressed a single predominant FAN identity, numerous cells within every cluster either expressed no FAN markers (6.1% of cells) or expressed markers for multiple FANs (Fig. 2B, Table S4). Overall, 46.6% of cells were cholinergic, 40.4% of cells were GABAergic, 30.1% of cells were glutamatergic, and 22.5% of cells expressed some combination of Glu, ACh, and GABA markers (Fig. 2B-2C, Table S4). However, these numbers likely overestimate the proportion of *Aedes* neurons capable of releasing multiple FANs. First, most putative multiple-FAN neurons express markers for different FANs at different levels; whether expression of lowly expressed secondary FANs is functionally relevant remains to be seen. Second, many multi-FAN neurons were classified as cholinergic based on ChAT expression alone (Fig. S4), and prior studies documented ChAT transcription without translation in many glutamatergic and GABAergic neurons (Lacin et al. 2019).

**Figure 2.**
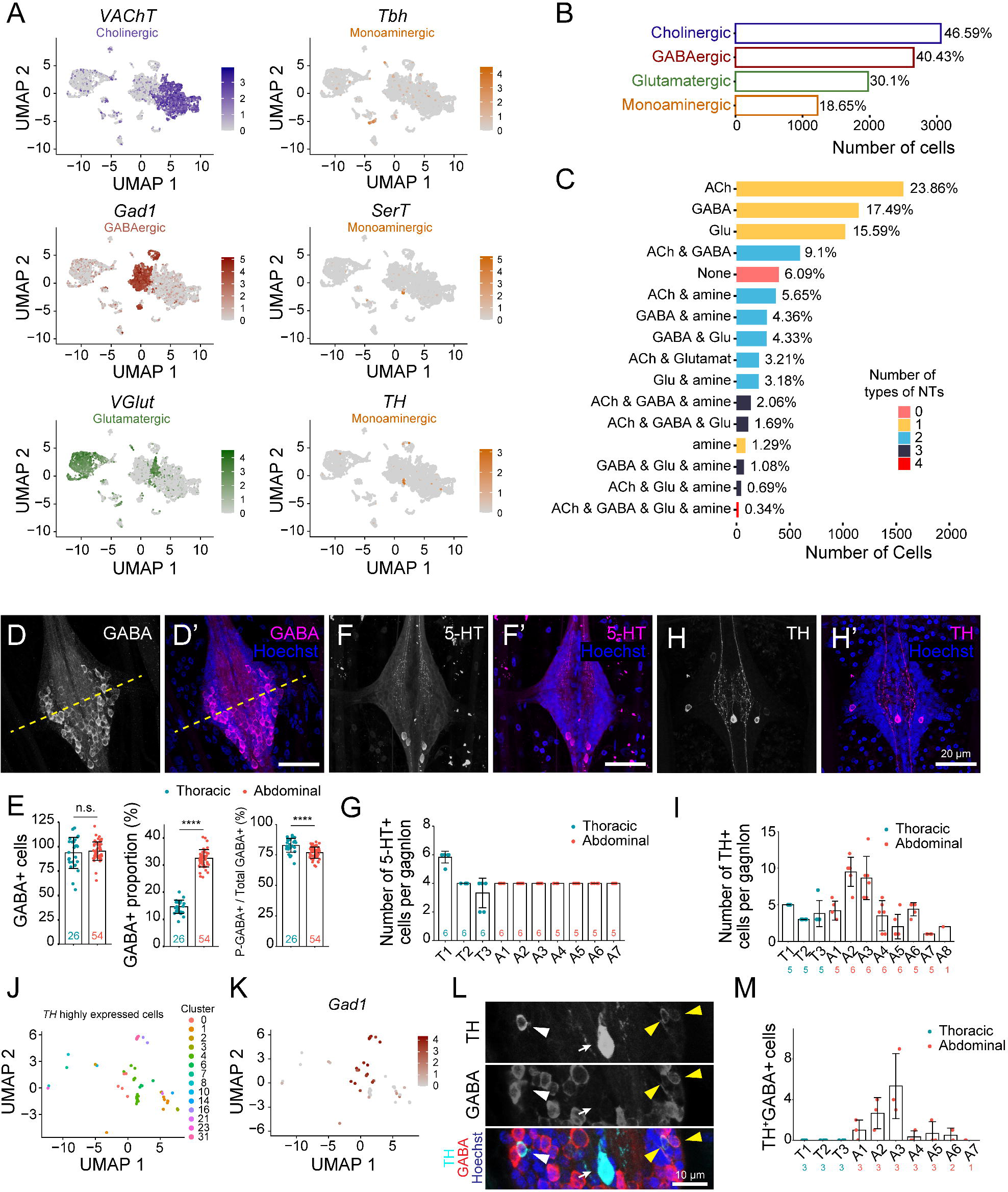
Neurotransmitter distribution in larval VNC cell types. (A) Feature plots showing expression of neurotransmitter marker genes *VAChT*, *Gad1*, *VGlut*, *Tbh*, *SerT*, and *TH*. (B) Histogram showing the percentage of neurons in the VNC cell atlas classified as cholinergic, GABAergic, glutamatergic, and monoaminergic according to marker gene expression. (C) Histogram depicting the frequency of different combinations of neurotransmitter co-expression. (D-I) *In vivo* expression patterns of neurotransmitters. (D) Maximum intensity projection image depicting the distribution of GABAergic neurons (labeled by anti-GABA immunoreactivity) in a representative abdominal segment. The yellow dashed line marks the midpoint of the ganglion along the AP axis; GABA+ cells principally accumulate in the posterior portion of each ganglion. (E) Quantification of the number of GABA+ cells, the proportion of cells in each ganglion that were GABA+, and the percentage of GABA+ cells located in the posterior half of the ganglion. Points represent measurements from an individual ganglion, and measurements were grouped according to segmental identity (thoracic or abdominal). *P < 0.05, unpaired t-test with Welch’s correction. (F) Maximum intensity projection image depicting the distribution of serotonergic neurons labeled by anti-5-HT immunoreactivity in a representative abdominal segment. (G) Quantification of 5-HT+ cell distribution in the VNC. T1 typically has 3 pairs of 5-HT+ cells and all other segments typically have 2 pairs of 5-HT+ cells. *P < 0.05, Kruskal-Wallis test with a post-hoc Dunn’s test. (H) Maximum intensity projection image depicting the distribution of dopaminergic neurons labeled by anti-TH immunoreactivity. (I) Quantification of cell distribution in the VNC. *P < 0.05, Kruskal-Wallis test with a post-hoc Dunn’s test. (J) UMAP plot of isolated, re-clustered *TH*+ cells shaded according to their cluster of origin. (K) Feature plots showing expression of GABAergic neuron marker genes *Gad1* and *VGAT* in *TH*+ cells. (L) Maximum intensity projection image depicting double-labeling of dopaminergic neurons with antibodies to TH and GABA. The white arrow indicates the medial high-TH neuron which is GABA-, the white arrowhead marks a TH+, GABA-cell, and yellow arrows indicate TH+, GABA+ double-positive cells. (M) Histogram depicting the number of TH+, GABA+ double-positive cells in each segment of the larval VNC. TH+, GABA+ double-positive cells are principally located in segments A2 and A3, which have the largest number of TH+ cells overall. *P < 0.05, Kruskal-Wallis test with a post-hoc Dunn’s test. Genotype for all panels: *brp-T2A-QF2w / +; QUAS-mcd8GFP / +*.

To explore patterns of FAN expression *in situ*, we used antibodies to GABA, ChAT, and Vglut to label larval VNCs. ChAT antibodies densely labeled structures in the neuropile but exhibited minimal labeling in the cortical cell layers (Fig. S4), and Vglut immunoreactivity was distributed across the entire ganglion, precluding unambiguous identification of glutamatergic and cholinergic neurons. In contrast, GABA immunoreactivity labeled a discrete subset of cells in each ganglion that were uniform in number and position (Fig. 2D-2E). Furthermore, thoracic and abdominal ganglia contained equivalent numbers of GABA-positive cells (thorax, 93.6 ± 16.0, n = 26; abdomen, 95.3 ± 9.5, n = 54), despite the fact that thoracic segments contained significantly more neurons and cells overall (Fig. 2E). In addition, GABA-positive cells were largely confined to the posterior compartment of each ganglion. A similar arrangement of inhibitory neurons has been described in locusts (Schmäh and Wolf 2003), however dipterans commonly have a network of distributed GABAergic neurons in each neuromere (Enell et al. 2007).

Next, we monitored the distribution of monoaminergic neurons and found that markers for monoaminergic transmitters were expressed in 18% of cells (Fig. 2B), corresponding to ∼450 cells per larval VNC assuming nearly uniform representation of each segment in our cell atlas. Despite this broad expression of monoaminergic markers, only ∼7% of monoaminergic cells (1.29% of cells overall) expressed monoamines as their exclusive transmitter, hence monoamines were predominantly co-expressed with one or more FANs (Fig. 2C). For example, we identified a single cluster of likely octopaminergic neurons that expressed monoamines as the predominant transmitter (cluster 26; 97.4% of cells expressed monoaminergic markers), and this cluster additionally expressed glutamatergic markers in >70% of cells (Table S3). For some monoaminergic types, we identified multiple subsets of cells that co-expressed distinctive FANs (Fig. S5). For example, we identified two additional populations of octopaminergic neurons: cluster 32, which expresses glutamatergic markers in 97.5% of cells, contains a subcluster that additionally expresses *Tbh* (47.5% of cells); and cluster 20, which expresses GABAergic markers in 93.75% of cells, contains a subcluster comprised of 29.5% of cells that express *Tbh* (Fig. S5, Table S4). Finally, we identified one cluster (cluster 4) that contained at least two distinct populations of monaminergic cells. Although cluster 4 primarily contains GABAergic cells (89.76% of cells express GABAergic markers), it contains two subclusters of GABA-negative monoaminergic cells including a population of dopaminergic cells that co-express *TH* and *Vglut* and an adjacent subcluster of serotonergic cells that express no FAN markers (Fig. 2A, S5). In total, we identified 6 clusters in which >30% of cells express monoaminergic markers (Table S3).

To characterize the distribution of monoaminergic neurons *in vivo*, we stained larval VNCs with antibodies against serotonin (5-HT) and TH to label serotonergic and dopaminergic neurons, respectively (Mysore et al. 2011; Vinauger et al. 2018). Our scRNA-seq studies identified one principal population of VNC serotonergic neurons within cluster 4 (Fig. 2A), and we found that 5-HT immunoreactivity is confined to pairs of bilaterally symmetric neurons in the larval VNC which are clustered with one another: three pairs in T1 and two pairs in other segments (Fig. 2F, 2G). We additionally detected punctate 5-HT immunoreactivity in the neuropile, particularly the anterior half of each ganglion, suggesting that 5-HT release may be concentrated in this region, and in projections to the periphery.

To identify potential regulators of serotonergic fate in the *Aedes* VNC, we isolated all serotonergic neurons from our dataset and queried for genes whose expression correlated with *SerT*. Among genes encoding annotated proteins, the four genes exhibiting the most highly correlated expression to *SerT* were involved in vesicular loading or biosynthesis of serotonin (Table S5), underscoring the validity of our approach. The next most highly correlated genes were transcription factors (TFs): the homeodomain TF *engrailed* (*en*) and the homeobox TF *Hox-c3a*. Expression of both TFs positively correlated with *SerT* expression, and using antibody staining we confirmed that en protein, which is expressed in only ∼20 cells per ganglion, is expressed in serotonergic neurons (Fig. S6). En is required for development and/or survival of serotonergic neurons *Drosophila* and mice (Lundell et al. 1996; Fox and Deneris 2012); our expression studies suggest that it may play similar roles in *Aedes* and that Hox-c3a may likewise promote serotonergic fate.

In contrast to 5-HT staining which labeled a stereotyped number of cells in each ganglion, TH staining revealed populations of cells in each ganglion that varied by number, TH intensity, soma position and projection patterns (Fig. 2H, 2I). First, while most ganglia contained 3-5 TH-positive cells, two abdominal ganglia (A2 and A3) contained 9-10 TH-positive cells on average. Second, TH staining intensity adopted a trimodal distribution: a single high-TH neuron was located medially, two medium-TH neurons were located posterior and laterally, and a variable number of low-TH neurons were distributed laterally (Fig. 2H, S7). TH is the rate-limiting enzyme in dopamine biosynthesis; these different populations may therefore have different capacities for dopamine signaling. Third, the high-TH neuron projected anteriorly and arborized throughout the neuropile whereas medium-TH neurons sent projections that entered the neuropil medially and largely arborized ipsilaterally; all TH-positive neurons appeared to project longitudinally between segments (Fig. 2H).

*TH* mRNA expression was distributed across multiple clusters in our scRNA-seq dataset (Fig. 2J), therefore we assayed for co-expression of *TH* and FAN markers to better define the different populations of dopaminergic neurons. We found that a substantial portion of *TH*-expressing neurons express *Gad1* and/or *GAT* and that *Gad1* and *TH* mRNA levels were inversely related (Fig. 2K, S7), hence low TH-expressing neurons are likely GABAergic and high TH-expressing neurons are likely GABA-negative. Indeed, double labeling of larval VNCs with antibodies to GABA and TH revealed co-expression in neurons located laterally, but not medially, within each ganglion (Fig. 2L). Finally, we found that these TH+ GABA+ double-positive neurons were most frequently observed in segments A2 and A3 (Fig. 2M), accounting for the overall increase in TH+ neurons in these segments (Fig. 2I).

### Identification of peptidergic neurons

In addition to classical neurotransmitters, neuropeptides comprise a diverse family of neuromodulators that serve as central regulators of larval physiology and behavior, controlling events including feeding, water balance, and ecdysis (Altstein and Nässel 2010). We next identified neuropeptide-expressing neurons in the larval VNC using two complimentary approaches: we assayed for expression of genes encoding neuropeptide precursor peptides, and we queried individual cluster markers for genes involved in neuropeptide release and for unannotated neuropeptides. We curated a list of 42 *Aedes* genes encoding neuropeptide precursors and identified 12 clusters that express a total of 18 neuropeptide genes using the following criteria: the neuropeptide gene is expressed by > 25% of cells in the cluster with an average scaled expression level > 1.0 (Fig. S8, Table S6). In addition, several neuropeptide genes including *ILPs*, *Limostatins*, and *sNPF* were highly expressed by small subsets of cells in some clusters (Fig. S8), expanding the likely repertoire of VNC-expressed neuropeptides. The larval VNC therefore appears to be a major source of peptidergic signals.

Many neuropeptides are expressed by neurons that lack FANs, and among seven clusters containing a large proportion (ranging from 19-60%) of neurons that expressed no FAN or monoamine marker genes, five of the clusters (cluster 19, 22, 24, 25, and 27) expressed neuropeptide genes (Fig. 3A, Table S3). Several observations suggest that these are bona fide peptidergic neurons. First, neuropeptide expression was both widespread within the clusters and highly restricted outside of these clusters (Table S7). Each of the clusters expressed a minimum of one neuropeptide gene in >80% of cells, and four of the clusters expressed neuropeptide genes in >95% of cells. Additionally, four of the clusters expressed multiple neuropeptide genes, including cluster 25 which co-expressed *glycoprotein hormone alpha2* (*Gpa2*) and *glycoprotein hormone beta5* (*Gbp5*) genes encoding the heterodimeric glycoprotein hormone thyrostimulin. Furthermore, expression of neuropeptide genes was restricted to a single cluster except for *allotropin*, which was expressed in both peptidergic cells (cluster 25) and cholinergic neurons (cluster 34). Second, in four of these clusters (19, 24, 25, and 27), >40% of cells lacked expression of any FAN or monaminergic marker genes, suggesting that they represent neurosecretory cells (Table S3). Three of these clusters additionally express genes encoding venom-like peptides including agatoxins (cluster 24, 26) and the turripeptide pal9.2 (cluster 22) whose function has not been characterized in mosquitos. Third, genes involved in neuropeptide processing including *peptidyl-α-hydroxyglycine-α-amidating lyase 2 (pal2)*, the prohormone convertase genes *amontillado* (*amon*) and *7b2*, and the metallocarboxypeptidase gene *silver* (*svr*) were identified as marker genes for these clusters (Fig. S8). We examined whether these peptidergic cells additionally expressed the bHLH transcription factor *dimmed*, which is associated with high levels of protein secretion in *Drosophila* (Hewes et al. 2003) but observed very little *dimmed* expression in any cluster. However, we identified *CrebA* as a marker for these peptidergic clusters (Fig. S9) and CrebA increases the secretory capacity of cells in some systems (Johnson et al. 2020).

**Figure 3.**
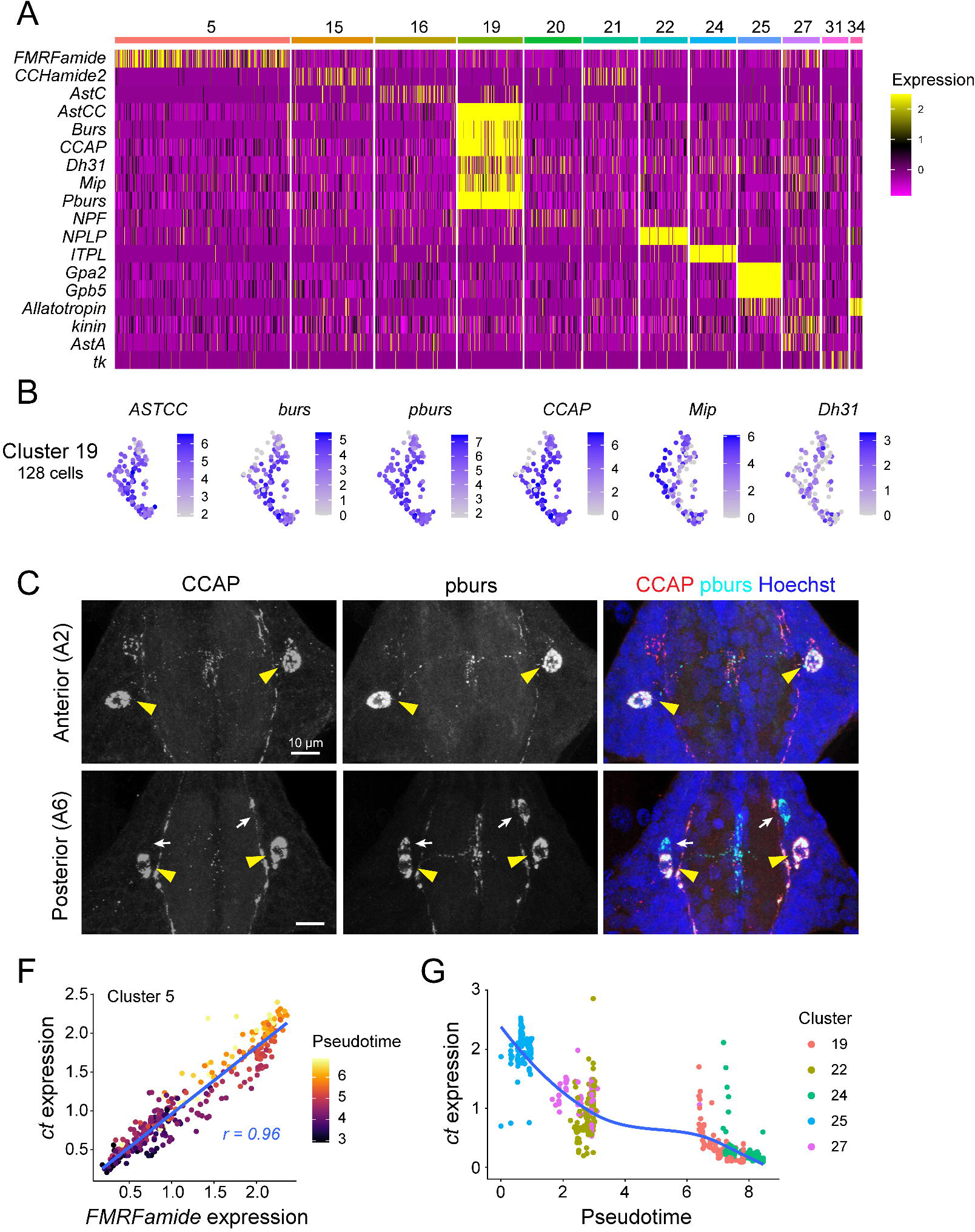
Identification of neuropeptide expressing cells. (A) Major neuropeptide-expressing cell clusters in the larval VNC. Heatmap depicts neuropeptide gene expression patterns in clusters that express neuropeptide genes in > 25% of cells at an average scaled expression of > 1.0. (B) Feature plots depicting expression of neuropeptide genes *ASTCC*, *burs*, *pburs*, *CCAP*, *Mip*, and *DH31* in cluster 19. Cells from this cluster co-express multiple neuropeptides. (C) Maximum intensity projections show Cluster 19 neurons identified *in situ* by immunostaining with antibodies to CCAP and pburs. All segments contain a bilaterally symmetric pair of neurons that co-express CCAP and pburs, and the posterior abdominal segment A6 but no other segment contains an additional pair of pburs-expressing cells that lack CCAP immunoreactivity. (F) FMRFamide expression is positively correlated with expression of the TF gene *ct*. Plot depicts expression levels of *FMRFamide* and *ct* in cluster 5. Dots represent individual cells, dot color represents pseudotime scores, and the linear regression line is shown in blue. (G) *ct* gene expression levels vary among different peptidergic clusters according to pseudo-time scores. *Ct* expression levels are plotted against pseudotime for the indicated peptidergic clusters. Dots represent individual cells and colors indicate cluster identity. Genotype for all panels: *brp-T2A-QF2w / +; QUAS-mcd8GFP / +*.

Among the neuropeptide-expressing clusters, we identified cluster 19 as a putative neurosecretory center that controls multiple elements of larval development. Cells in this cluster express neuropeptides including *allatostatin CC* (*AstCC*), *bursicon* (*burs*), *partner of bursicon* (*pburs*), *crustacean cardioactive peptide* (*CCAP*), *diuretic hormone 31* (*DH31*), and *Myoinhibiting peptide precursor* (*Mip*) that play essential roles in controlling ecdysis behavior, diuresis, and feeding in a broad range of insects and crustaceans (Loveall and Deitcher 2010;

Honegger et al. 2011; Lahr et al. 2012; Verbakel et al. 2021; Zieger et al. 2021). All the cells in this cluster expressed multiple neuropeptides: half of the cells expressed all six of these neuropeptide genes simultaneously, 31.3% co-expressed 5 neuropeptide genes, and 17.2% co-expressed 4 neuropeptides genes (Fig. 3B, S10). To localize these cells *in vivo* we stained larval VNCs with antibodies to pburs and CCAP, which were among the most broadly expressed of the six neuropeptides (100% of cluster 19 cells express *pburs*, 93.8% express *CCAP*, Fig. S10). These antibodies both labeled one pair of bilaterally symmetrical neurons in each ganglion that extend processes within the ganglion and out to the periphery (Fig. 3C). Outside of the soma, CCAP and pburs exhibited a punctate distribution in the neuropil that was not completely overlapping, suggesting that they may be released in non-overlapping domains. We identified an additional pair of pburs-positive cells confined to posterior abdominal ganglia (A6, A7, A8) which lacked CCAP immunoreactivity, and processes of these neurons appeared to innervate distinct regions of the neuropil from the CCAP-pburs co-expressing cells (Fig. 3C). Taken together with our scRNA seq data indicating that other neuropeptide genes, *Mip* and *DH31*, are expressed in cluster 19 subsets, these results suggest that cluster 19 comprises morphologically and functionally distinct sub-groups of peptidergic neurons.

Neuropeptides are known to be co-expressed with FANs (Croset et al. 2018), and indeed we identified six clusters in which cells co-expressed a single predominant FAN and a neuropeptide: *FMRFa* in glutamatergic neurons (cluster 5), *NPF* and *AstC* in GABAergic neurons (clusters 10 and 16), and *RYamide*, *CCHa2*, and *allotropin* in cholinergic neurons (cluster 1, 21, and 34) (Fig. 3A). Although orthologous neuropeptides are co-expressed with FANs in the *Drosophila* adult VNC (Allen et al. 2020), we identified two principal differences between neuropeptide-FAN co-expression in *Drosophila* and *Aedes*. First, some *Drosophila* cells co-express more than one neuropeptide together with a FAN, but neuropeptide-FAN co-expression in *Aedes* occurred exclusively in cells expressing a single FAN. We did, however, identify one cluster that co-expressed a FAN, monoamines, and a putative neuropeptide (cluster 26). Second, neuropeptide-FAN pairing in *Drosophila* and *Aedes* matched only in the case of *AstC*, which is expressed in GABAergic neurons in both systems (Brunet Avalos et al. 2019), suggestive of stage and/or organism specializations for function of these neuropeptides.

Compared to peptidergic clusters that largely lacked FANs (cluster 19, 22, 24, 25, and 27), neuropeptide expression was more variable in clusters that co-expressed FANs (cluster 1, 5, 10, 21, and 34) (Fig. 3A). To explore the transcriptional basis of this variability in neuropeptide gene expression, we focused on the largest such cluster, which expressed *FMRFamide* in 56.2% of cells (cluster 5, Fig. 3A). Correlation analysis revealed that expression of the TF *cut* (*ct*) was highly correlated with *FMRFamide* expression in cluster 5 cells, and we additionally found that *ct* and *FMRFamide* expression increased over pseudotime (Fig. 3F). Likewise, we found that levels of *ct* varied according to pseudotime across several other peptidergic clusters (Fig. 3G), suggesting that *ct* is a candidate temporal TF that regulates peptidergic neuron differentiation. Consistent with this possibility, we found that Ct protein accumulates to varying degrees in different cells within the VNC (Fig. S11), and *ct* expression likewise varies according to developmental progression in an identified neuroblast lineage in *Drosophila* (Seroka et al. 2022).

### Identification of cell cluster markers

Overall, neuropeptide genes were among the most highly enriched markers for individual clusters in our entire dataset (Table S7). In contrast, patterns of neurotransmitter expression provided a coarse functional map of VNC cell types. To facilitate more detailed analyses of the developmental origin and precise function of each cell cluster in the *Aedes* VNC we identified marker genes that were both broadly expressed and enriched in each cluster using differential expression analysis (Table S7). This analysis yielded an average of 185 markers per cluster (6,487 markers total), however many genes appeared as markers for multiple clusters. For example, *Vglut* and *VAChT* were identified as marker genes for each of the glutamatergic and cholinergic clusters, respectively, and both *GAD* and *VGAT* were identified as marker genes for GABAergic neurons. Neurotransmitter receptors likewise appeared as markers for most clusters, and although certain neurotransmitter receptors were enriched in a small number of clusters, most were broadly expressed (Fig. S12). In contrast, we found that individual monoamine and neuropeptide receptors were often highly enriched in one or a small number of cell clusters (Fig. S12). For example, the *Aedes* T*achykinin receptor TkR99D* and *sNPF receptor* genes were principally expressed in cluster 25, which additionally expressed the *Myosuppressin receptor*. Likewise, the *Ecdysis triggering hormone receptor* gene and the *CCHamide2 receptor* gene were principally expressed in subsets of cluster 19 and 34 cells, respectively. Hence, these peptidergic populations are likely subject to peptidergic modulation. As with cluster 25, several clusters expressed more than one neuropeptide receptor gene, hence patterns of neuropeptide receptor co-expression mark subsets of cells within individual clusters. Finally, we found that in contrast to other neuropeptide genes which showed restricted expression domains, the *Dopamine/ecdysteroid receptor (DopEcR)* gene, which encodes a G-protein coupled dual receptor for dopamine and ecdysteroids (Srivastava et al. 2005), was expressed in most cell clusters. The dual ligand capacity of DopEcR suggests that it may function as a coincidence detector, and its broad expression suggests that it may broadly regulate VNC neurons, for example in responses to stressors as in *Drosophila* (Petruccelli et al. 2020) or priming cells for pupation as in Lepidoptera (Kang et al. 2019).

The most differentially expressed genes in the *Drosophila* larval nervous system are TFs (Li et al. 2020; Seroka et al. 2022), and indeed we found that most clusters expressed a unique repertoire of TFs (Figure S9). First, we identified TFs that mark individual clusters, but this was principally true for peptidergic cells, possibly reflecting an increased expression distance of peptidergic cells from other clusters. For example, we identified *odd-paired* (*opa*) as a marker for peptidergic cluster 19 and *eyes absent* (*eya*) as a marker for peptidergic cluster 24; both of these TFs mark peptidergic cells in the *Drosophila* VNC, suggesting that these factors may play conserved roles in peptidergic fate (Miguel-Aliaga et al. 2004; Simon et al. 2019). More commonly, individual TFs were markers for multiple clusters, including cases in which a TF marked multiple clusters with a particular NT identity (e.g., *acj-6* and *unc-4*, which mark cholinergic clusters) and clusters with distinct NT identities (e.g., *kn* is highly expressed in cholinergic clusters 11, 15, and peptidergic cluster 24). Finally, we also found that numerous TFs were highly expressed in subsets of neurons within individual clusters, likely reflecting developmentally and/or functionally distinct subsets of neurons within the cluster.

### Ventral ganglia positions are defined by Hox gene expression along the A-P axis

In both invertebrates and vertebrates, patterns of Hox gene TF expression specify segment identity along the anterior-posterior (A/P) axis (McGinnis and Krumlauf 1992). Our anatomical studies demonstrated that thoracic and abdominal ganglia differ in cell number and composition, therefore we examined the distribution of Hox gene expression across our VNC cell atlas to determine whether any cell clusters are restricted to thoracic or abdominal ganglia. For these studies we monitored expression of the *Antennapedia* (*Antp*), *Ultrabithorax* (*Ubx*), *abdominal-A* (*abd-A*), and *Abdominal-B* (*Abd-B*), which are expressed in partially overlapping domains along the AP axis (Fig. S13A). First, we used antibodies to validate the AP distribution of these proteins in *Aedes* larvae. As anticipated, Antp protein is largely restricted to the thoracic neuromere: >90% of cells in thoracic ganglia and <5% of cells in abdominal ganglia expressed Antp (Fig. S13B). Although monoclonal antibodies that selectively recognized *Drosophila* Ubx, Abd-A and Abd-B exhibited no immunoreactivity in the *Aedes* VNC, an antibody that recognizes both Ubx and Abd-A labeled no cells in T1 and >90% of cells in T2 and all posterior segments. In our scRNA cell atlas we found that mRNA for each of these Hox genes was broadly expressed: we identified *Antp*, *Ubx*, *Abd-A* and/or *Abd-B* expression in 87.3% of cells (Fig. S13C). Furthermore, although every cluster contained cells that expressed each of *Antp*, *Ubx*, *Abd-A,* and *Abd-B*, some clusters were enriched for a particular Hox gene, suggestive of enrichment of these clusters at particular A/P positions in the VNC (Fig. S13D). Finally, we note that *Abd-B* expression was largely absent from three clusters (25, 32, 34), however this may be a product of the small size of these clusters and restricted expression domain of *Abd-B*. Overall, these results suggest that ganglia in each segment contain cells from each cluster. However, we note that cells expressing each Hox gene were spatially segregated within some clusters, for example *Antp* and *Abd-A* expression in cluster 11, suggesting that distinct thoracic and abdominal sub-clusters of some neuronal types likely exist.

### Identification of immature neurons in mosquito ventral ganglia

Our analysis of neurotransmitter and neuropeptide expression revealed that 7/35 clusters contained a large proportion of cells (ranging from 19-60%) expressing no FAN or monoamine marker genes. Among these, we identified cluster 10 as a likely source of neural precursors and differentiating neurons. First, cluster 10 contained comparable proportions of cholinergic, GABAergic, glutamatergic and to a lesser degree monoaminergic cells (33.83%, 25.87%, 24.88%, and 14.93%, respectively). Overall, >77% of cluster 10 cells expressed one or no neurotransmitter, hence the cluster was comprised of a heterogeneous mixture of neurotransmitter fates (Table S4). Second, we examined the identify of cluster 10 marker genes and found that many of them are involved in *Drosophila* neural stem cell proliferation and differentiation (Fig. 4A, Table S7). These markers include *headcase* (*hdc*), which functions as negative regulator of neurogenesis (Maierbrugger et al. 2020) and is a marker gene for undifferentiated neurons (Brunet Avalos et al. 2019; Dillon et al. 2022); the tumor suppressor *brain tumor* (*brat*) and RNA binding protein *Syncrip* (*Syp*) which regulate neuroblast proliferation and differentiation (Lee et al. 2006; Yang et al. 2017); and *ciboulet (cib*) and *Ten-Eleven Translocation* (*Tet*), which are markers for immature adult neurons (Croset et al. 2018; Corrales et al. 2022). Third, we calculated cell cycle phase scores based on expression of canonical marker genes and found that most cells in cluster 10 (62.7%) but no other cluster expressed G2/M markers, indicative of recent or pending cell division (Fig. 4B). Fourth, genes involved in neurotransmitter biosynthesis and transport were expressed to varying degrees in cluster 10 cells, suggestive of a continuum of differentiation. We therefore hypothesized that this cluster contains neural precursors and/or immature neurons undergoing differentiation.

**Figure 4.**
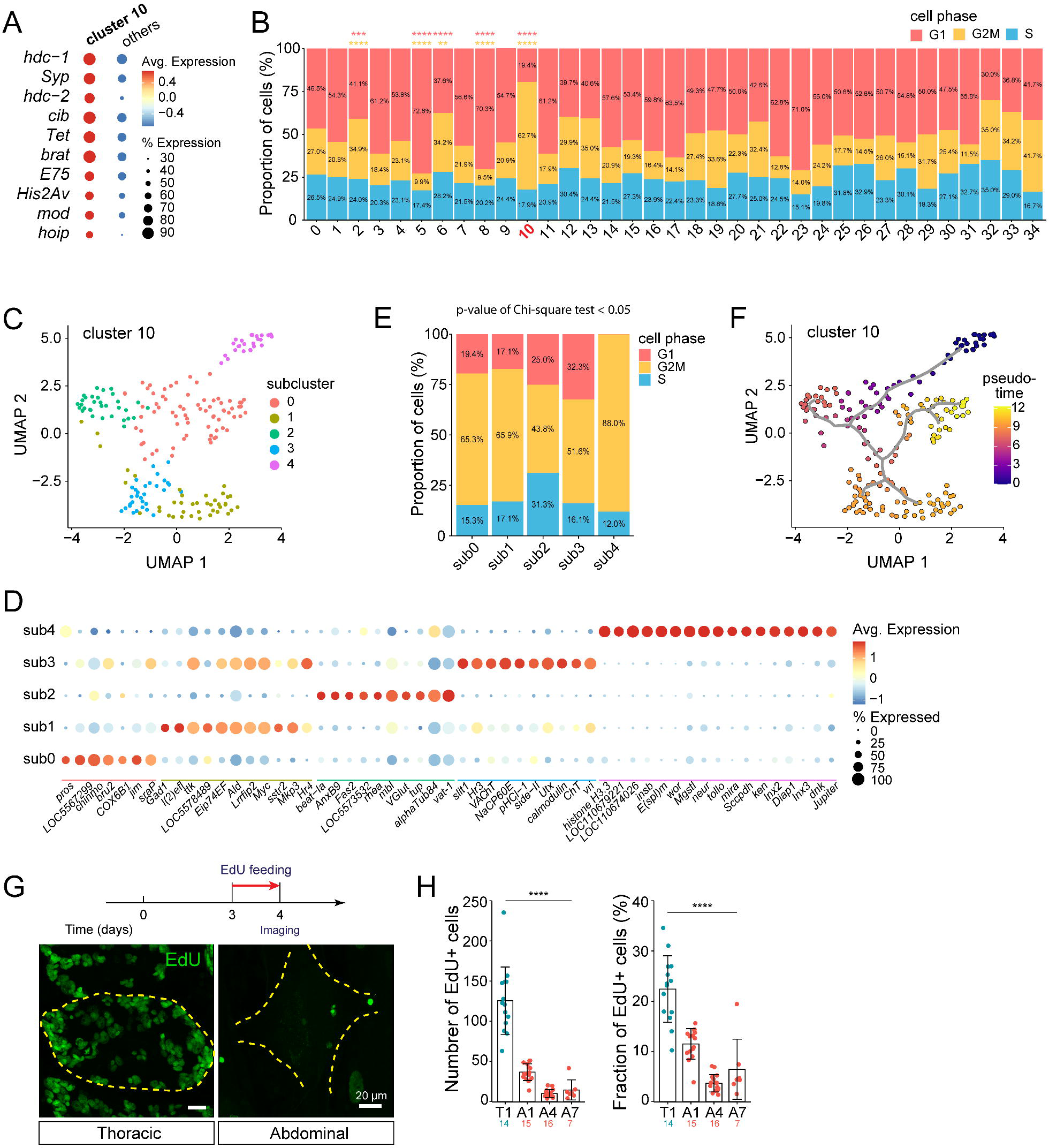
Identification of neural progenitors in the VNC. (A) Dot plot showing expression of cluster 10 marker genes within cluster 10 and across all other clusters. (B) Cell-cycle phase analysis showing the percentage of cells in G1, G2/M, and S phase, assigned according to marker gene expression. *P<0.05, Pearson’s Chi-square test with a post-hoc Bonferroni Adjustment for multiple comparisons. (C-F) Identification of neuroblasts, intermediate progenitors, and differentiating neurons. (C) Seurat UMAP plot depicting 5 cluster 10 sub-clusters. (D) Dot plot depicting marker gene expression for cluster 10 sub-clusters. Neurotransmitter marker genes were highlighted by bold font. (E) Cell-cycle phase analysis of cluster 10 sub-clusters. Sub-cluster 4 had a significantly larger portion of cells in G2M phase than other clusters and had no cells expressing G1 markers. *P<0.05, Pearson’s Chi-square test with a post-hoc Bonferroni Adjustment for multiple comparisons. (F) Pseudo-time trajectory analysis of neural progenitor cells from cluster 10, anchored at sub-cluster 4. Cells are colored according to a lookup table that depicts their pseudo-time score along the trajectory, with the darkest color indicating the most-immature neurons. (G) Identification of mitotic populations in the VNC. *Top*, schematic depicting the workflow for EdU incorporation experiments. Bottom, representative maximum intensity projection images depict EdU immunoreactivity in thoracic and abdominal ganglia. (H) Plots depict the number (left) and fraction of cells in the indicated segments (T1, A1, A4, A7) which incorporated EdU. Boxes represent mean values, bars represent standard deviation, and points represent measurements from individual segments. *P < 0.05, Kruskal-Wallis test with a post-hoc Dunn’s test. Genotype: wild type in (G, H); all other panels: *brp-T2A-QF2w / +; QUAS-mcd8GFP / +*.

To further characterize the cellular composition of this putative neuronal precursor cluster, we isolated cluster 10 from all other cells in the dataset and subjected it to independent clustering analysis. We found that cluster 10 contains five sub-clusters, each of which was enriched in a set of marker genes suggestive of different stages in neuronal differentiation (Fig. 4C, Table S8). First, sub-cluster 4 expressed canonical neuroblast marker genes including *miranda* (*mira*), *insensible* (*insb*), *worniu* (*wor*), and *neuralized* (*neur*) (Boulianne et al. 1991; Ikeshima-Kataoka et al. 1997; Schuldt et al. 1998; Carney et al. 2012; Komori et al. 2018), suggesting that sub-cluster 4 is comprised of neuroblasts (Fig. 4D). Consistent with this possibility, we found that sub-cluster 4 was principally populated by cells in G2/M phase (Fig. 4E). Additionally, cells from sub-cluster 4 expressed lower levels of mature neuronal marker genes including *Brp*, *nSyb*, *Syt*, and *CadN* than other cluster 10 sub-clusters or a global average of *Aedes* VNC neurons (Fig. S14). Second, sub-cluster 0 contained a heterogeneous cell population that expressed genes involved in neuroblast differentiation including the TFs *prospero* (*pros*) and *chronologically inappropriate morphogenesis* (*chinmo*) as well as markers for immature neurons including *hdc*, the zinc finger TF *jim*, and the actin-binding factor ciboulet (*cib*) (Corrales et al. 2022), suggesting that sub-cluster 0 contains products of neuroblast differentiation (Fig. 4D). In addition, the anterior Hox gene *Antp* was enriched in sub-cluster 0, suggesting that these cells are enriched in the thoracic neuromere (Table S8). Antp was likewise expressed by a larger proportion of cells in other cluster 10 sub-clusters than in the dataset overall, suggesting that cluster 10 may be enriched in the thorax. Third, sub-clusters 1, 2, and 3 expressed GABAergic, glutamatergic, and cholinergic markers, respectively, as well as markers for immature neurons, suggesting that these clusters contain differentiating neurons adopting distinct FAN cell fates (Fig. 4D).

To probe the relationships between cluster 10 sub-clusters, we performed single cell trajectory analysis using Monocle (Trapnell et al. 2014; Qiu et al. 2017; Cacchiarelli et al. 2018). Specifically, we mapped trajectories through pseudotime to infer developmental relationships between the clusters by anchoring the putative neuroblasts – cells from sub-cluster 4 – at the origin of the trajectory. Consistent with our inferences based on marker gene expression, we found that cells followed a trajectory from sub-cluster 4 through a portion of sub-cluster 0 and into branches comprised of sub-clusters 1, 2, 3 and the remainder of sub-cluster 0, indicative of a differentiation axis from top to bottom in UMAP space (Fig. 4F). This trajectory suggests that sub-cluster 0 contains cells at different developmental stages, and indeed sub-cluster 0 cells in the distal portion of the pseudotime trajectory expressed markers for differentiating neurons including FANs (Table S8). We extended this single cell trajectory analysis to include all cells (Fig. S15), and this analysis revealed two additional features in the dataset. First, we found that one additional cluster (cluster 6) exhibited a low pseudotime value comparable to cluster 10 (Fig. S15), expressed markers of immature neurons including *Lim1*, *Rbp6*, and *scylla* (Corrales et al. 2022), and lacked a single predominant FAN identity (Tables S3, S6). Hence, cluster 6 may represent another population of neurons undergoing differentiation. Second, different clusters that shared a given FAN identity could be distinguished by their pseudotime scores (Fig. S15), which may provide an entry point to define the developmental connections between the different clusters; we discuss one factor that covaries with pseudotime within a particular lineage below.

To identify cluster 10 cells *in situ*, we stained larval *Aedes* VNCs with antibodies against *Drosophila* versions of the neuroblast marker *mira* (sub-cluster 4) and immature neuron markers *hdc* and *pros* (sub-cluster 0), however none of the antibodies yielded specific cross-reactivity (Table S1). We therefore assayed for mitotic activity as a proxy for neuroblast division. First, we stained VNCs with antibodies to phosphorylated histone H3 (PH3) to label mitotic cells in L2 and L4 larvae. We found mitotic activity in the VNC was substantially higher in thoracic than abdominal ganglia, and principally found in L2 VNCs: (L2 thorax, 35.7 ± 6.7 cells, n = 18; abdomen, 2.9 ± 2.6, n = 37) (Fig. S16). To confirm these results, we fed L2 larvae with the thymidine analog EdU and monitored EdU incorporation 24 h later (Fig. 4G). Consistent with our PH3 staining results, we observed substantially more labeling in thoracic than abdominal ganglia (T1, 125.8 ± 42.0 cells, n = 14; A1, 36.7 ± 10.5 cells, n = 15 larvae) (Fig. 4H). Taken together, these results suggest that post-embryonic neuroblast divisions principally occur in thoracic segments during early larval stages.

### Larval neurons persist in the adult stage

Our studies of mitotic activity in the in the larval VNC suggest that *Aedes* larvae exhibit minimal cell addition within abdominal ganglia during larval development. We therefore hypothesized that a large proportion of larval neurons persist in the adult mosquito nervous system, and we examined this possibility in the adult abdominal VNC. We found that the cellular composition of abdominal ganglia in *Aedes* larvae and adults was remarkably similar, with no discernable difference in the number or distribution of cells (Fig. 5A, 5B). Of note, the average number of neurons present in each ganglion was constant from Day 3 of larval development through adulthood (Fig. 5B). Furthermore, markers of neuronal subsets exhibited similar distributions in larvae and adults: GABAergic neurons exhibited comparable posterior enrichment in larvae and adults, and both serotonergic and dopaminergic neurons were present in corresponding numbers and positions in larvae and adults (Fig. 5C-5E). To assess rates of cell turnover more broadly, we assayed for EdU incorporation as a proxy for cell division and AnnexinV staining to label apopotic cells during pupal stages. Although we saw extensive EdU incorporation outside of the nervous system, we saw no signs of EdU labeling in abdominal ganglia (Fig. 5G, 5H). Likewise, when we used TUNEL to identify apoptotic cells we observed minimal labeling in abdominal ganglia of larvae or post-eclosion adults (Fig. 5I, 5J, S17), supporting the notion that adult ganglia are largely populated by persistent larval neurons.

**Figure 5.**
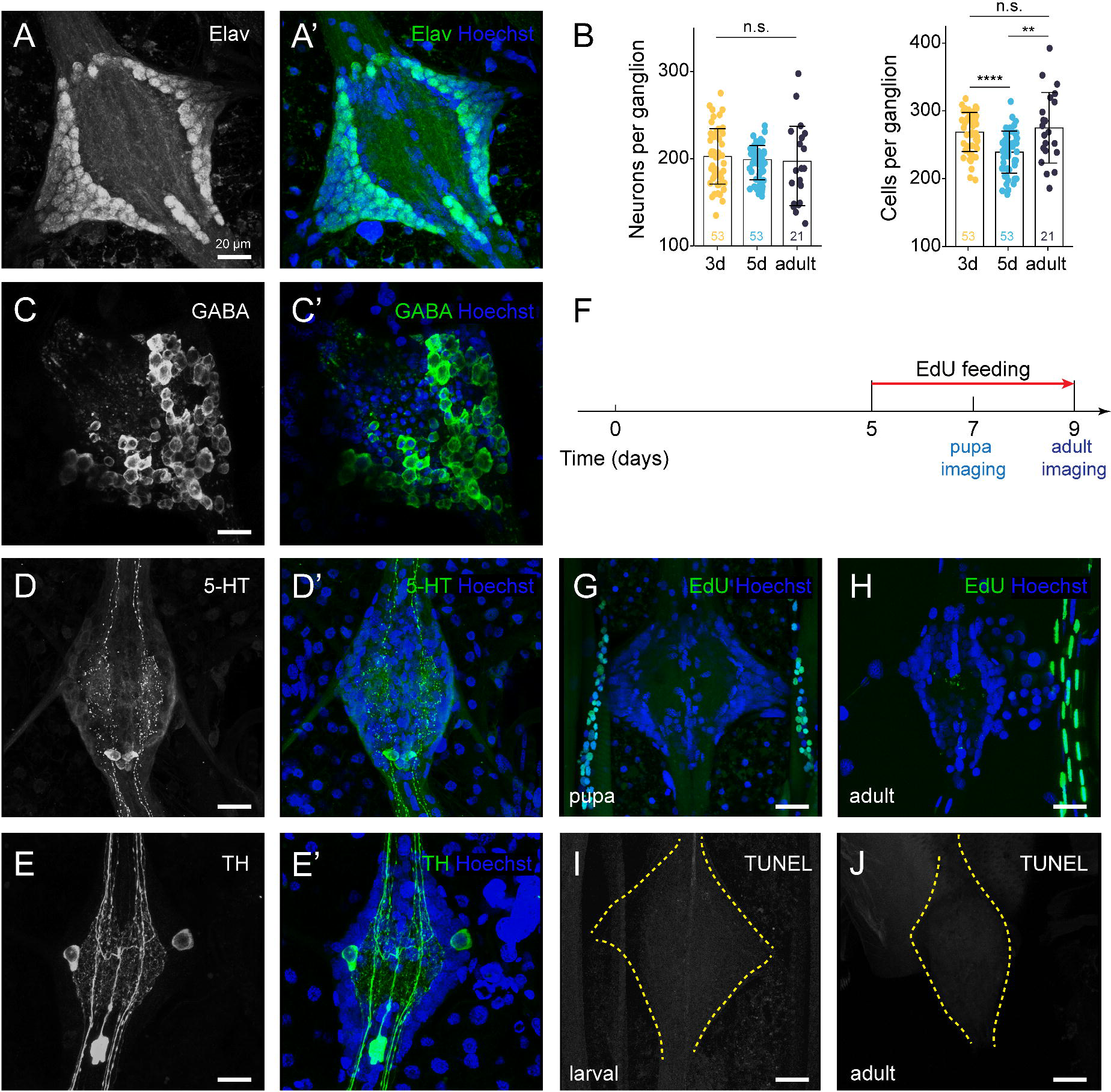
Neuronal identities in *Aedes* adult ventral ganglia. (A) Maximum intensity projections showing representative abdominal ganglia from segment A6 of 2-day post-eclosion *brp-T2A-QF2w / +; QUAS-mcd8GFP / +* adults stained with antibodies to Elav to label neurons and Hoechst to label all nuclei. (B) Quantification of neuronal number and total cell number across larval and adult development (3 days after larval hatching, 5 days after larval hatching, 2 days after adult eclosion). Bars depict mean values, points represent measurements from individual ganglia, and error bars represent standard deviation. N.s., not significant, one-way ANOVA analysis. (C-E) Distribution of GABAergic, serotonergic and dopaminergic neurons in adult abdominal VNC. Maximum intensity projection images depict the distribution of (C) GABAergic neurons (labeled by anti-GABA immunoreactivity), (D) serotonergic neurons labeled by anti-5HT immunoreactivity, and (E) dopaminergic neurons labeled by anti-TH immunoreactivity in segment A6 of 2-day post-eclosion adults. (F) Schematic workflow schedule of EdU feeding and imaging. Animals were continuously maintained in EdU-containing water from Day 5 of larval development. (G-H) Representative images depicting distribution of anti-EdU immunoreactivity in abdominal ganglia from pupa (G) and 2-day post-eclosion adult (H). Although EdU incorporation was not readily detectable in abdominal ganglia, other tissues including tracheal cells (adjacent to ganglia) were readily labeled. (I-J) Representative images depict TUNEL labeling of the larval (I) and adult abdominal ganglion (J). TUNEL labeling revealed very few apoptotic cells at either developmental stage. Figure S17 depicts positive controls for labeling efficacy. Genotype: wild type in (G, H); all other panels: *brp-T2A-QF2w / +; QUAS-mcd8GFP / +*.

## Discussion

Several recent studies have explored the cellular makeup of the adult mosquito nervous system (Severo et al. 2018; Raddi et al. 2020; Kwon et al. 2021; Cui et al. 2022; Herre et al. 2022), however other life history stages have been less extensively studied. Here, we explored neuroanatomy and cellular diversity in the larval *Aedes* VNC. First, we find that the basic structure of the VNC is preserved through larval development and into adulthood, particularly in abdominal segments. Second, we generated a single cell transcriptional atlas that reveals neuronal diversity in the *Aedes* VNC (Fig. 6); we additionally identify markers for putative glial populations. Taken together, these datasets provide the identity of cell types that likely control key elements of larval behavior and physiology and markers that will facilitate comprehensive functional studies in the VNC. Third, we identify likely progenitor cells and juvenile neurons, which represent the building blocks of the adult nervous system. Finally, we demonstrate that abdominal segments of the adult VNC exhibit remarkable similarity to larval counterparts and are likely composed primarily of persistent larval neurons, underscoring the importance of understanding larval VNC development and function.

**Figure 6.**
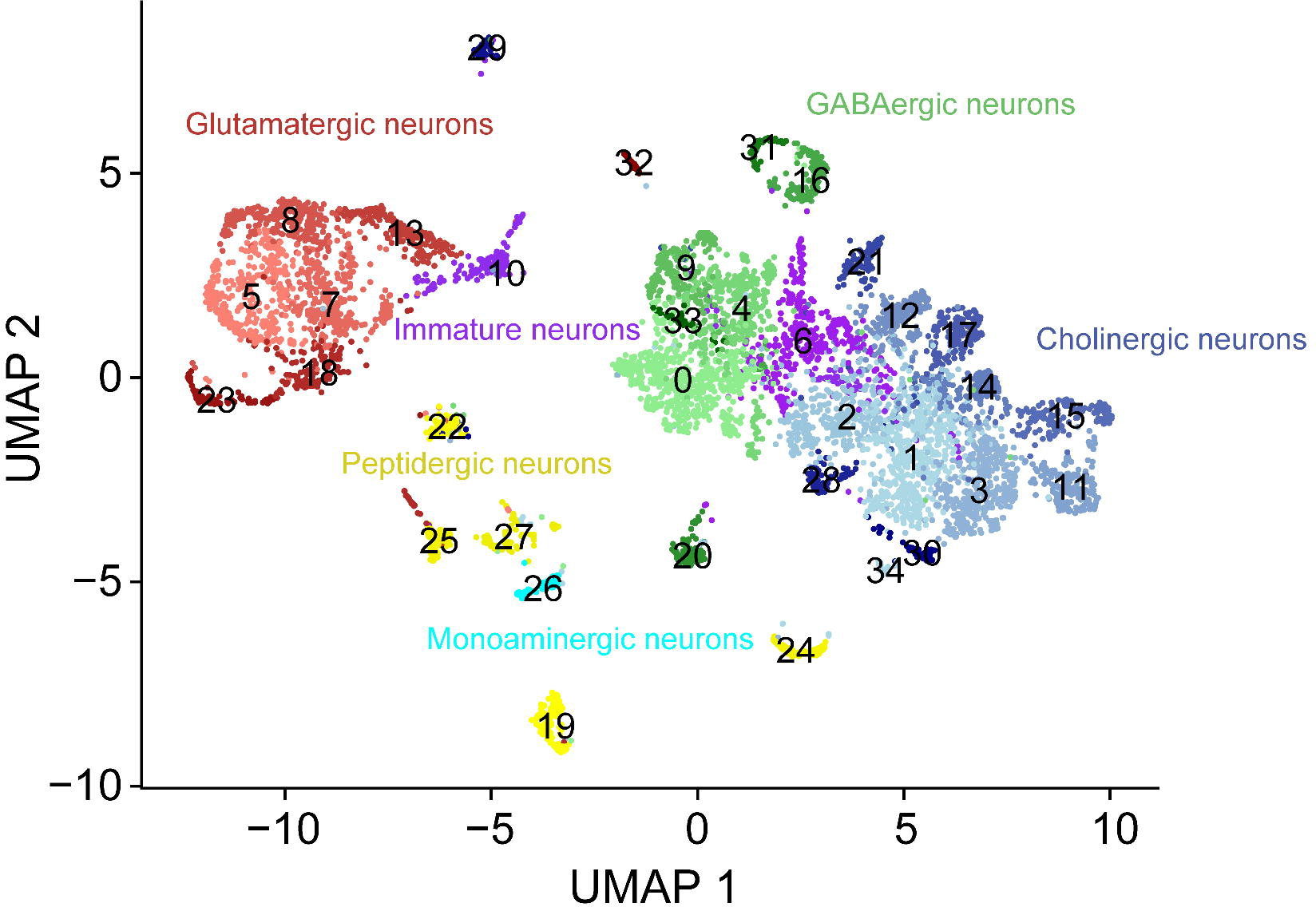
Annotated *Aedes* larval VNC cell atlas. The UMAP plot depicts 6,564 neurons grouped into 35 clusters. Immature neurons (cluters 6 and 10), and all other clusters are color-coded according to predominant transmitter type.

In contrast to the *Aedes* larval VNC, the cellular repertoire of the *Drosophila* VNC has been systematically characterized. As a result, the *Drosophila* VNC has served as an invaluable template to guide study fundamental properties of other insect nervous systems, including *Aedes*. However, comparative genomics suggests that *Aedes* and *Drosophila* share a common ancestor from ∼250 million years ago (Hedges et al. 2006; Arensburger et al. 2010), and the nervous systems of the two organisms have diverged extensively (Smarandache-Wellmann 2016), hence the utility of comparative approaches is limited for many aspects of *Aedes* biology. First, although the VNCs share common design elements including the segmental organization of bilaterally symmetrical ganglia, placement of central neuropil and longitudinal tracts, the VNC morphology is distinct, with fused neuromeres in *Drosophila* and a chain of segmental abdominal ganglia in *Aedes*. Furthermore, the *Drosophila* VNC contains twice as many cells during the larval stage as *Aedes* (Scott et al. 2001; Birkholz et al. 2015; Lacin et al. 2019) and the degree of correspondence in cell types populating the VNCs is not known. Our study points towards extensive functional divergence: with only one exception, we found that neuropeptides and FANs were expressed in different combinations. Finally, although the *Aedes* and *Drosophila* VNCs subserve similar overall functions, they drive different locomotion strategies, receive inputs from different sensory organs, and control different appendages.

Overall, our study reveals the identity of 35 clusters of VNC cells that differ according to their expression landscape. As with other similar studies, these clusters likely represent many more cell types, and this point is illustrated by our targeted analysis of monoaminergic and peptidergic cells. For example, we identified serotonergic neurons as a discrete subset of a primarily GABAergic cell cluster (cluster 4); unlike other cells in the cluster, these serotonergic neurons expressed neither GABAergic markers nor markers for other FANs. We likewise identified several populations of dopaminergic neurons which were themselves discrete subsets of larger cell clusters, and our anatomical studies revealed the presence of at least three distinct dopaminergic sub-populations which differed in position, number, and TH expression. Hence, while cluster identity is a useful starting point for identifying related groups of cells, targeted analysis of cluster markers will likely reveal a larger number of functionally related subsets.

One approach to mosquito population control involves targeting larval populations with treatments referred to as larvicides that block growth and/or maturation of larvae, often targeting juvenile hormone signaling (Tsang et al. 1988). While effective, many currently used larvicides including methoprene are toxic to other marine life including some fishes and aquatic arthropods (Lawler 2017; Peterson and Rolston 2022). Our studies identify components of at least two axes of neuropeptide signaling that could be leveraged for population control. First, we identified neurons that express neuropeptides and neuropeptide receptors with established roles in feeding and growth including NPF, CCHamide2, allostatins, and allotropin. The distribution of neuropeptide receptors is suggestive of extensive crosstalk among these peptidergic neurons: for example, the DH31 receptor is expressed in cells that express CCHamide2, CCHamide2 receptor is expression in cells that express allotropin, and the allotropin receptor is in turn expressed in a subset of CCHamide2-expressing cells. It therefore seems likely that treatments targeting this entire peptidergic network would have profound effects on larval growth. Second, we identified one cluster of neurons that we hypothesize functions as a neurosecretory center controlling ecdysis; these cells represent an additional target for population control. Insect ecdysis, particularly adult eclosion, and post-ecdysis events including wing expansion are coordinated by a neuropeptide cascade that initiates appropriate motor programs: peripheral release of ecdysis triggering hormone (ETH) commits animals to ecdysis, with a central cascade of ecdysis hormone (EH) and CCAP driving ecdysis behaviors and burs driving post-ecdysis behaviors including wing expansion (White and Ewer 2014). Our studies suggest that cluster 19 neurons are at the center of *Aedes* eclosion; these neurons express the ETH receptor, which plays a role in activating burs-expressing neurons in *Drosophila* (Kim et al. 2006) as well as CCAP and burs. Our studies likewise identified putative motor neuron targets of these neuropeptides: the glutamatergic clusters 19 and 32 express CCAP receptor and burs receptor (Rickets), respectively.

One surprising takeaway from our studies is the remarkable similarity in the cellular composition of larval and adult abdominal ganglia. Adult brain development in dipterans and other insects that undergo complete metamorphosis involves extensive remodeling of the embryonic/larval brain, including removal of cells by programmed cell death, addition of cells produced during post-embryonic divisions of glial and neuronal progenitors, and widespread remodeling of larval circuitry during metamorphosis (Levine et al. 1995; Tissot and Stocker 2000). While we see evidence for extensive post-embryonic neurogenesis in thoracic ganglia (Fig. 4G-4H), the situation is different in the abdomen. Cell numbers in abdominal ganglia are constant from early larval stages through early adulthood, and unlike thoracic ganglia we see limited mitotic activity in larval or pupal abdominal ganglia. Moreover, we see no evidence of widespread cell death in larval or pupal abdominal ganglia. Instead, we find that identified classes of larval neurons are present in the same number and position in adults, and additionally have comparable projection patterns. Furthermore, our analysis of likely neural progenitor cells suggests that immature neurons are enriched in thoracic segments.

In addition to the limited turnover of larval neurons, developmental timing likely provides an additional constraint to large-scale reorganization of larval circuitry during *Aedes* metamorphosis. Remodeling of *Drosophila* larval mushroom body neurons and peripheral sensory neurons during metamorphosis requires 18 h or more for pruning events alone, followed by multiple days for regrowth of new processes (Yaniv and Schuldiner 2016). In addition, *Aedes* pupae are motile and therefore maintain functional sensorimotor circuits while incorporating adult-specific circuit elements. Altogether, these observations support the notion that abdominal ganglia in adults are populated largely by persistent larval neurons, further underscoring the value of understanding larval nervous system development.

Several questions arise from our observations. First, how extensively are larval neurons and circuits remodeled to give rise to adult circuits? We defined a panel of antibodies that label structural elements in the VNC including major axon fascicles, synapses, and cell bodies, and demonstrated that antibodies to specific neurotransmitters and neuropeptides label projections in discrete subsets of cells. If combined with super resolution imaging approaches, these markers should facilitate fine-scale morphological analysis of dopaminergic, serotonergic, and some peptidergic neurons in larvae and adults. More broadly, the cluster markers we identified should facilitate development of transgenes for sparse labeling of neurons of interest. Second, to what extent are functions of larval neurons preserved in the adult VNC? Although many spared neurons are rewired in *Drosophila*, some retain larval patterns of connectivity in adults. For example, the moonwalker descending neuron (MDN) elicits backward locomotion in larvae and adults, and following remodeling during metamorphosis, MDN connectivity with the Pair1 interneuron is re-established in adults (Lee and Doe 2021). As discussed above, the short duration of metamorphosis likely constrains the extent of nervous system remodeling, so it will be particularly interesting to explore larval functions of persistent adult neurons that are involved in pathogen transmission and host interactions. For example, nerve cord ligation studies suggest that the abdominal ganglion controls blood meal size: adults with severed nerve cords exhibit hyperphagia and abdominal rupture from overfeeding (Gwadz 1969). Given the well-documented roles for serotonergic neurons in controlling feeding behaviors (Kinney et al. 2014; Tierney 2020), VNC serotonergic neurons are one candidate neural substrate for this behavior. We identified a single principal population of serotonergic neurons in our cell atlas and found that both the number and position of serotonergic neurons are preserved from larvae to adults. It remains to be determined how extensively these neurons remodel their connectivity and whether they likewise contribute to satiety signaling in larvae.

## Materials and Methods

### Mosquito husbandry

*Ae. aegypti* wild-type Liverpool IB12 (LVP) and *brp-T2A-QF2w; QUAS-mcd8GFP* transgenic strains were maintained and reared at 28 °C with a photoperiod of 14h light / 10h dark cycle. Hatched larvae were maintained in plastic trays filled with 1L distilled water and fed with tropical fish food.

### Sample preparation for single cell RNA-seq

Protocols for cell isolation and library preparation were adapted from protocols developed for analysis of the *Drosophila* larval VNC (Nguyen et al. 2021). 5-day old *Ae. aegypti* larvae were sorted under a fluorescent stereomicroscope and incubated on ice to immobilize larvae prior to dissection. Larvae were pinned dorsal side up on sylgard dishes (Dow corning) containing ice-cold PBS, filleted along the dorsal midline, and VNCs were isolated from surrounding tissue and transferred to transferred to 1.5ml Eppendorf tubes containing dissociation buffer (450 µl PBS supplemented with 1% BSA solution and 1 mg/mL collagenase type X). A total of 50 VNCs were dissociated (15-20 VNCs in each tube) at 37 °C with mechanical agitation (mixing at 1000 rpm with manual trituration every 10 min). Dissociated cell suspensions were filtered through 70 µm nylon filters to remove cell debris, stained with propidium iodide (PI) to label dead cells, and GFP positive/PI negative neurons were captured using a FACSArial II (BD Biosciences, San Jose, CA). 61,646 single cells were collected and resuspended in 50 µl PBS + 1% BSA solution after centrifuging for 10 mins at 300x g and 4 °C.

### Library construction and sequencing

Library construction was performed using a Chromium Single Cell 3’ v3.1 kit (10X Genomics). The single cell suspension was diluted to ∼1000 cells/µl and loaded onto the chip to target a recovery of approximately 10,000 cells. Following RT-PCR, GEMs were stored at 4 °C overnight followed by cDNA amplification, fragmentation, and indexing the next day. Library quality was evaluated on an Agilent TapeStation (size range of reads from 300 bp to 800 bp with an average size of 500 bp). Sequencing was performed on an Illumina NovaSeq SP 100 Cycle Flow Cell by the Northwest Genomics Center, Seattle.

### Gene lists and Annotation

We combined information from Ensembl BioMart (https://metazoa.ensembl.org/index.html), Vectorbase (vectorbase.org), and Flybase (flybase.org) into an integrated pipeline for orthology calls. Our list of annotations includes 20128 genes in total, 12458 with an NCBI gene description annotation, and 9319 with *Drosophila* orthologues.

### Computational analysis for single cell transcriptomic atlas

#### Mapping and counting

A customized mosquito reference for mapping was generated using the CellRanger (version 6.1.1) ‘mkref’ function, and 86.2% of reads could be mapped to the *Ae. aegypti* genome (AaegL5.0, GCF_002204515.2 on NCBI) after sequencing. A 10X Genomics expression matrix with 362,382,783 reads in total and 27,030 average reads per cell was obtained after mapping and counting by CellRanger (version 6.1.1).

#### Quality Control

The following analyses were performed on R and RStudio, following quality control parameters used in a comparable dataset from *Drosophila* larvae (Vicidomini et al. 2021). For quality control cells that had fewer than 500 genes, 1000 UMIs, and log (genes) / log (UMIs) ratios of less than 0.8 were removed. Cells with a mitochondrial gene proportion greater than 18%, ribosomal gene proportion less than 5% or greater than 40%, and heat shock gene proportion greater than 5% were removed. We also removed genes with counts less than 5 and expressed in less than 2 cells. Cell multiplets were identified by DoubletFinder (version 2.0.3) and removed.

#### Data analysis with Seurat

7,965 single cells with 12,680 genes expressed were kept as the original dataset. Seurat package (version 4.3.0) in R was used for the following analyses. 2,000 highly variable features were identified through the ‘FindVariableFeatures’ function and the dataset was scaled by the ‘ScaleData’ function. To determine the dimensionality of the dataset, we first performed Principal Component Analysis (PCA) and then used the Elbow-Plot and JackStraw-Plot tests together with an evaluation of PC-heatmaps for selecting proper principal components (PCs). 35 PCs were considered to explain the variance of the dataset. Non-linear dimensional reduction was performed by using the ‘RunUMAP’ function and followed by finding clusters through the ‘FindClusters’ function with the resolution set at 0.8. The clustering result was visualized in the UMAP plot with 25 clusters (cluster 0 to cluster 24). Average number of total UMIs, number of genes, and sample sizes were calculated at the cluster level. Clusters with lower numbers of total UMIs, number of genes, and sample sizes were considered as heterogenous clusters. Expression levels of neuronal and glial marker genes were checked in the dataset to identify non-neuronal clusters. After removing heterogenous and non-neuronal cells, the filtered dataset contained 6,564 single cells.

The same analysis pipeline was used for the filtered dataset. 33 PCs were considered to explain the variance of the dataset. Non-linear dimensional reduction was performed by using the ‘RunUMAP’ function and followed by finding clusters through the ‘FindClusters’ function with the resolution set at 1.5. The clustering result was visualized in the UMAP plot with 35 clusters (cluster 0 to cluster 34). Markers of each cluster were defined using the ‘FindAllMarkers’ function by selecting genes positively expressed in the cluster with the minimum percentage set to 0.5 and log fold change threshold as 0.32. In this way, we identified 2,231 genes as cluster markers for this atlas and used them to annotate the identities of clusters.

### Sub-clustering analysis

#### Immature neurons

Immature neurons from cluster 10 were subsetted from the filtered dataset, yielding a new dataset consisting of 201 single cells. The new dataset was reanalyzed and subclustered using the workflow described above. In this case, 500 genes were identified as highly variable features, 8 PCs were selected to generate the UMAP plot with a resolution of 1.0. Marker genes for each subcluster were defined using the method described above.

#### Dopaminergic neurons

Dopaminergic neurons were subsetted by selecting neurons with normalized expression of the *TH* gene at a level greater than 1, which yielded a new dataset consisting of 60 single cells. These cells were subsequently analyzed for expression of FAN marker genes.

#### Serotonergic neurons

Serotonergic neurons were subsetted by selecting neurons with *SerT* gene normalized expression greater than 1, yielding a new dataset consisting of 118 single cells. These cells were subsequently analyzed using correlation analysis as described below.

#### Peptidergic cluster 19

Neurons from peptidergic cluster 19 were subsetted from the filtered dataset, yielding a new dataset consisting of 128 single cells. These cells were subsequently analyzed for expression of several different neuropeptide genes.

### Trajectory analysis

Pseudotime analysis was conducted for neurons from cluster 10 and all neurons in the dataset, respectively, using a Monocle3 analysis pipeline (https://github.com/cole-trapnell-lab/monocle-release/tree/monocle3_alpha). Seurat objects were converted into the Monocle3 cell_data_set objects using the ‘as.cell_data_set’ function in SeuratWrappers package. The pseudotime score for each single cell was calculated using the ‘learn_graph’ function. The starting point for the analysis of neurons from cluster 10 was assigned as sub-cluster 4, and the start point for the analysis in the whole dataset was assigned as cluster 10.

### Cell-cycle scoring

A list of marker genes for S phase and G2 and M (G2/M) phases was generated for this analysis by identifying *Ae. aegypti* orthologs of *Drosophila* cell phase marker genes used in previous research (Tirosh et al. 2016; Jevitt et al. 2020). A cell cycle score was calculated for each single cell by the ‘CellCycleScoring’ function from Seurat R package. Cells lacking detectable expression level of marker genes for S and G2/M were assigned as G1 phase. Each cell was assigned to a cell cycle phase and the percentage of cells at each cell phase was calculated at the cluster level or subcluster level.

### Correlation analysis

To identify gene interactions at the single cell level, we used Markov Affinity-based Graph Imputation of Cells (MAGIC) algorithms for imputing missing values and restoring the structure of the data. Imputation of the Seurat object was performed by the ‘magic’ function with default parameters. The R package ‘dismay’ (https://github.com/skinnider/dismay) was used to calculate correlations in the gene co-expression networks after imputation. Gene interactions were visualized by ‘ggplot’ using the expression levels in each single cell after MAGIC imputation.

### Immunostaining

*Ae. aegypti* larvae were pinned dorsal side up on sylgard dishes (Dow corning) containing ice-cold PBS, filleted along the dorsal midline, and pinned open to expose the VNC. Intestines, fat bodies, and trachea were removed, samples were rinsed in PBS, and subsequently fixed for 30 min in PBS + 4% formaldehyde (PBSFA) on a rotary shaker. Following fixation, samples were washed twice with PBS for 2 min each and subsequently permeabilized PBST (PBS + 0.3% Triton X-100) for 15 min (3 washes, 5 min each). Specimens were blocked for 15 min in PBST containing 5% NGS (supplier) and incubated in primary antibody diluted in blocking buffer at 4 °C overnight. The following day, samples were washed twice with PBS for 2 min each, three times with PBST for 10 min each, and incubated with the secondary antibody and Hoechst (1:500) at room temperature for 1-2 hours in blocking buffer. Following staining, samples were washed with 800 µl PBST 3 times for 10 min each at room temperature, rinsed twice in PBS, and mounted in Fluoromount G mounting medium (Electron Microscopy Sciences). Slides were dried at room temperature for a minimum of 1 hr prior to imaging. Details on antibody sources and dilutions are provided in Table S1.

### TH immunofluorescence intensity measurements

Fluorescence intensity of TH immunoreactivity was measured in maximum intensity projections of confocal stacks using ImageJ (Schneider et al. 2012). Corrected total cell fluorescence (CTCF) was calculated by using the following formula: CTCF = Integrated Density – (Area of selected cell × Mean fluorescence of background readings), with regions adjacent to the cell lacking immunoreactivity measured as the background. Based on the distribution of CTCF values cells were identified as low-, medium-, or high-TH-expressing cells: low, anti-TH CTCF values below 15,000; medium, anti-TH CTCF values above 15,000 but below 50,000; high, cells with anti-TH CTCF values above 50,000.

### EdU incorporation

Larvae were transferred to basins containing water supplemented with EdU to a final concentration of 0.2 mM and fed for the indicated time. Dissected larvae in PBS, fixed and blocked samples as described above in immunostaining method. Prepared the Click-iT reaction mix as instructed by the manufacturer. Added 500 µl of Click-iT reaction mix and incubated with rocking for 30 min at room temperature. Washed with PBST 3 times, 5 min each time. Added primary antibody at the blocking buffer and processed for immunostaining as described above.

### TUNEL Labeling

Apoptotic cells were identified via terminal deoxynucleotidyl transferase dUTP nick end labeling (TUNEL) with the In Situ Cell Death Detection Kit, Fluorescein (Roche, 11684795910), according to a protocol adapted from use in *Drosophila* tissue (Chimata et al. 2022). Larval and adult specimens were collected and dissected as described above. After fixing with PBSFA, samples were washed with PBST three times for 10 min each. Samples were incubated in freshly prepared Sodium Citrate buffer (495 µL of 100 mM Sodium Citrate + 5 µL of 10% Triton X-100) for 30 min in dark at 65 °C. Following washing in PBST (three times for 10 min each), samples were incubated in 100 µL TUNEL reaction mixture for 1 hr at 37 °C in a humidified chamber in the dark. After labeling, samples were washed three times in PBST for 10 min each, with Hoechst added to the last wash for nucleic acid staining. As positive control, a subset of samples was incubated with 250 U/mL DNase (Invitrogen) in PBS for 10 min at room temperature prior to TUNEL labeling. As a negative control, a subset of samples was incubated in 100 µL of Label Solution without terminal transferase.

## Data and code availability

Raw sequencing files (fastq) and digital expression matrices are available from the Gene Expression Omnibus under accession number PRJNA1008733. Code used in this analysis is available from GitHub.

## Acknowledgements

We are grateful for feedback on the manuscript from members of the Parrish laboratory. We thank the following members of the *Drosophila* community for generously sharing antibodies: Benjamin Altenheim, Clemens Cabernard, Claude Desplan, Aaron DiAntonio, Kirsten Guss, Valerie Hilgers, Adrian Moore, James Skeath, and Benjamin White; antibodies were additionally obtained from the Developmental Studies Hybridoma bank, created by the NICHD of the NIH and maintained at The University of Iowa. We also thank the McBride laboratory for sharing mosquitos, Jessica Huang, Kevin Huynh, and Ashley Pham for assistance with dissections; Jessica Huang for assisnance with library preparation. This work was supported by an award from the Royalty Research Foundation at the University of Washington and a grant from the National Institutes of Health to the MBL Neurobiology course MBL (R25NS063307).

## Author contributions

Conceptualization (T. Morita, J.Z. Parrish, C. Yin), data acquisition (C.Y.), data curation (C.Y.), formal analysis (C.Y.), funding acquisition (J.Z. Parrish), project administration and supervision (J.Z. Parrish), resources (T. Morita, J.Z. Parrish), software (C. Yin), data visualization (C. Yin), writing–original draft (J.Z. Parrish, C. Yin), writing–review and editing (T. Morita, J.Z. Parrish, C. Yin).

## Supplemental Figures

**Figure S1.** Quality control measurements for larval *Aedes* VNC dataset. (A) Histogram showing the log_10_ distribution of cells expressing the indicated number of UMIs. Red line represents the nUMI cutoff, with cells below 1,000 UMI excluded from further analysis. (B) Histogram showing the log_10_ distribution of cells with the indicated number of expressed genes (nGene). Red line represents the nGene cutoff; cells with fewer than 500 expressed genes were excluded from further analysis. (C) Histogram showing the distribution of cells with indicated proportion of mitochondrial transcripts. Red line represents the mitochondrial proportion cutoff (18%), above which cells were eliminated from further analysis. (D) Histogram showing the distribution of cells with the indicated proportion of transcripts from ribosomal genes. Cells with ribosomal gene proportions less than 5% or greater than 40% were eliminated from further analysis. Genotype: *brp-T2A-QF2w / +; QUAS-mcd8GFP / +*.

**Figure S2.** Identification of putative glial cells in the larval *Aedes* VNC. (A) Cell atlas from initial clustering, which contains 25 distinct cell clusters. Putative glial cells, identified based on marker gene expression, are indicated. (B) Cells from this putative glial cluster were isolated and reclustered to reveal putative glial subtypes. Feature plots depict expression of glial marker genes including the pan-glial marker *repo*, cortex glia marker *wrapper*, surface glia marker *gemini* and three astrocyte glial markers *wun2, Eaat1* and *Gat*. Genotype: *brp-T2A-QF2w / +; QUAS-mcd8GFP / +*.

**Figure S3.** Distribution of neuronal marker gene expression in the *Aedes* larval VNC cell atlas. Feature plots depict expression of *nSyb*, *Syt*, and *CadN* which are present in all clusters. Genotype: *brp-T2A-QF2w / +; QUAS-mcd8GFP / +*.

**Figure S4.** Neurotransmitter marker gene expression in the larval *Aedes* VNC. (A-D) Feature plots showing expression of neurotransmitter marker genes *ChAT*, *VGAT*, *Vmat*, and *Dat*. (E) Histogram showing number and percentage of cells expressing the indicated neurotransmitter marker genes. (F) Venn diagram showing the number of cells expressing the cholinergic marker genes *VAChT* and *ChAT*. Among the 329 ChAT+ VAChT-cells, 240 expressed an additional FAN marker gene (*VGlut*, *VGAT* and/or *Gad1*). (G) Maximum projections of confocal stacks depicting anti-ChAT immunostaining of the abdominal ventral ganglion, which was additionally labeled with the nucleic acid stain Hoechst. ChAT immunoreactivity is distributed throughout the neuropil but largely absent from the cortical cell layers. Genotype: *brp-T2A-QF2w / +; QUAS-mcd8GFP / +*.

**Figure S5.** Identifying subsets of monoaminergic neurons. Neurons that highly expressed monoaminergic marker genes were isolated from the VNC cell atlas and re-clustered to identify monoaminergic subsets. (A-D) Feature plots depict expression of the monoaminergic marker genes *Tbh*, *SerT*, *Dat*, and *TH*. Note that these genes are expressed in mutually exclusive sub-populations. (E) Seurat UMAP plot depicting monoaminergic neuron subtypes assigned basis on marker gene expression. Genotype: *brp-T2A-QF2w / +; QUAS-mcd8GFP / +*.

**Figure S6.** Identification of transcriptional markers for serotonergic neurons. Neurons that expressed the serotonergic marker *SerT* were isolated and re-analyzed. Feature plots depict expression of *SerT* (A) and two transcription factor genes *en* (B) and *Hox-C3a* (C) whose expression is highly correlated with *SerT* (D-E). Expression correlation plots additionally depict *Gad1* expression colored according to a lookup table. Note that cells with the highest levels of *SerT*, *en*, and *Hox-C3a* exhibit low expression of *Gad1*. (F) Maximum projections of confocal stacks depict representative images of the abdominal VNC labeled with antibodies to serotonin (5-HT) and en as well as the nucleic acid stain Hoechst. Cells co-expressing 5-HT and en are indicated with yellow arrowheads. Genotype: *brp-T2A-QF2w / +; QUAS-mcd8GFP / +*.

**Figure S7.** Classification of dopaminergic neurons according to TH levels and GABA expression. (A) Histogram showing the distribution of dopaminergic cells with indicated intensities of anti-TH fluorescence (CTCF, corrected total cell fluorescence). Cells were classified according to breaks in the distribution: low TH cells had TH levels below 15,000 CTCF; high TH cells had TH levels greater than 50,000 CTCF; medium TH cells had intermediate TH levels (between 15,000 and 50,000 CTCF). (B) Histogram depicts the average number of low-, medium-, and high-expressing TH cells in each segment of the larval VNC. N=6 animals, 251 neurons. (C-E) Relationship between TH and GABA expression. (C) Histogram showing the number of GABA-positive and GABA-negative low-, medium-, and high-expressing TH cells. Most TH+, GABA+ double-positive cells have low anti-TH levels. N = 3 animals, 129 neurons. (D) High *TH*-expressing cells (n = 60) were isolated from the global VNC dataset and reclustered. The feature plot depicts expression of the GABAergic neuron marker gene *VGAT* in dopaminergic neurons. (E) *TH* and *Gad1* expression levels are inversely related in dopaminergic neurons. Scatter plot depicts expression of TH (x-axis), Gad1 (y-axis), and VGAT (shading) in 106 dopaminergic neurons. High-TH-expressing cells express low levels of *Gad1* and *VGAT1*, and vice-versa. Genotype: *brp-T2A-QF2w / +; QUAS-mcd8GFP / +*.

**Figure S8.** Expression patterns of neuropeptides and peptide processing enzymes. Dot plots depict expression levels of (A) neuropeptide genes and (B) neuropeptide processing enzyme genes in each cell cluster of the VNC cell atlas. Dot color indicates mean expression level across the cluster and dot diameter indicates the proportion of cells in the cluster with nonzero expression of the gene. Genotype: *brp-T2A-QF2w / +; QUAS-mcd8GFP / +*.

**Figure S9.** Transcriptional markers of VNC cell clusters. Dot plots depict expression of transcription factors identified as marker genes for the indicated cell clusters, grouped according to transmitter identity: cholinergic (A), GABAergic (B), aminergic and peptidergic (C), and glutamatergic neurons (D). Genotype: *brp-T2A-QF2w / +; QUAS-mcd8GFP / +*.

**Figure S10.** Co-expression of neuropeptide genes in neurosecretory cluster 19. (A) Histogram showing the number and proportion of cluster 19 cells that express the following neuropeptide genes: *DH31*, *Mip*, *burs*, *CCAP*, *pburs*, and *ASTCC*. (B) Histogram showing the number and proportion of cluster 19 cells that co-express multiple neuropeptides in the same cell. Half of cluster 19 cells express 6 neuropeptide genes simultaneously. Genotype: *brp-T2A-QF2w / +; QUAS-mcd8GFP / +*.

**Figure S11.** Expression pattern of the transcription factor gene *ct*. (A) Feature plot showing *ct* expression across the VNC cell atlas. (B) Maximum projections of confocal stacks depicting anti-Ct immunoreactivity in a representative segment of the abdominal VNC additionally labeled with the nuclear dye Hoechst. As with *ct* mRNA, Ct protein is broadly expressed at varying levels in the VNC. Genotype: *brp-T2A-QF2w / +; QUAS-mcd8GFP / +*.

**Figure S12.** Expression patterns of neurotransmitter and neuropeptide receptor genes. Dot plots depict expression levels of (A) FAN receptor genes, (B) monoamine receptor genes, and (C) neuropeptide receptor genes in each cell cluster of the *Aedes* VNC cell atlas. Genotype: *brp-T2A-QF2w / +; QUAS-mcd8GFP / +*.

**Figure S13.** Expression patterns of Hox transcription factors. (A) Schematic diagram indicating putative expression patterns of Hox genes mapped alongside a composite image of the larval VNC and segmental nerves visualized in a fillet preparation of a *brp-T2A-QF2w, QUAS-mcd8GFP* larva. (B) Maximum intensity projections depict representative results from antibody staining of thoracic (T2) and abdominal (A4) segments of the larval VNC with antibodies to Antp and Ubx/Abd-A (antibodies recognize both Ubx and Abd-A). (C) Feature plots depicting Hox gene expression across the larval VNC cell atlas. (D-E) Histograms depict (D) the overall number of cells and (E) the proportion of cells from each cluster which express the indicated Hox genes. Colors are used to indicate neuronal subtypes. Genotype: *brp-T2A-QF2w / +; QUAS-mcd8GFP / +*.

**Figure S14.** Marker gene expression in neural progenitors and immature neurons. Dot plot depicts expression levels of pan-neuronal and FAN marker genes in cluster 10 sub-clusters. Putative neuroblasts (sub-cluster 4) exhibit limited expression of neuronal marker genes, sub-cluster 0 expresses neuronal markers but no FAN markers, and sub-clusters 1-3 expressed neuronal markers and an individual FAN marker. Genotype: *brp-T2A-QF2w / +; QUAS-mcd8GFP / +*.

**Figure S15.** Global pseudotime analysis of VNC development. (A) Pseudo-time trajectory analysis of the entire cell atlas, anchored at cluster 10. Cells are colored according to their pseudo-time score along the trajectory, with darker colors marking more immature neurons and lighter colors indicating less immature neurons. (B) UMAP plot highlighting cluster 6 and cluster 10, both of which are comprised principally of immature neurons. (C) Box plots showing pseudotime scores for each cluster. Clusters are grouped according to their principle FAN and ordered according to pseudotime scores. Genotype: *brp-T2A-QF2w / +; QUAS-mcd8GFP / +*.

**Figure S16.** Identification of mitotic cells in the larval VNC. (A) Maximum intensity projection of confocal stack depicting anti-PH3 immunoreactivity in thoracic ventral ganglion additionally labeled with anti-Lamin antibodies to facilitate cell counting. (B) Quantification of PH3+ cells in thoracic and abdominal segments at day 3 and day 5 of larval development. *P < 0.05, Kruskal-Wallis test with a post-hoc Dunn’s test. Ganglia number (N) for each sample: 3-day thorax (18), 3-day abdomen (37), 5-day thorax (29), 5-day abdomen (75). Genotype: wild type.

**Figure S17.** Positive control for efficacy of TUNEL labeling. Larval tissue was fixed, permeabilized, and pre-incubated with the endonuclease DNaseI for 10 min to induce DNA fragmentation prior to TUNEL labeling. Images depict TUNEL labeling of a larval ganglion (A) additionally labeled with Hoechst (B).

