## Supplementary figures and images for "A cell atlas of the larval *Aedes aegypti* ventral nerve cord"

### Figure S1_ver5.png

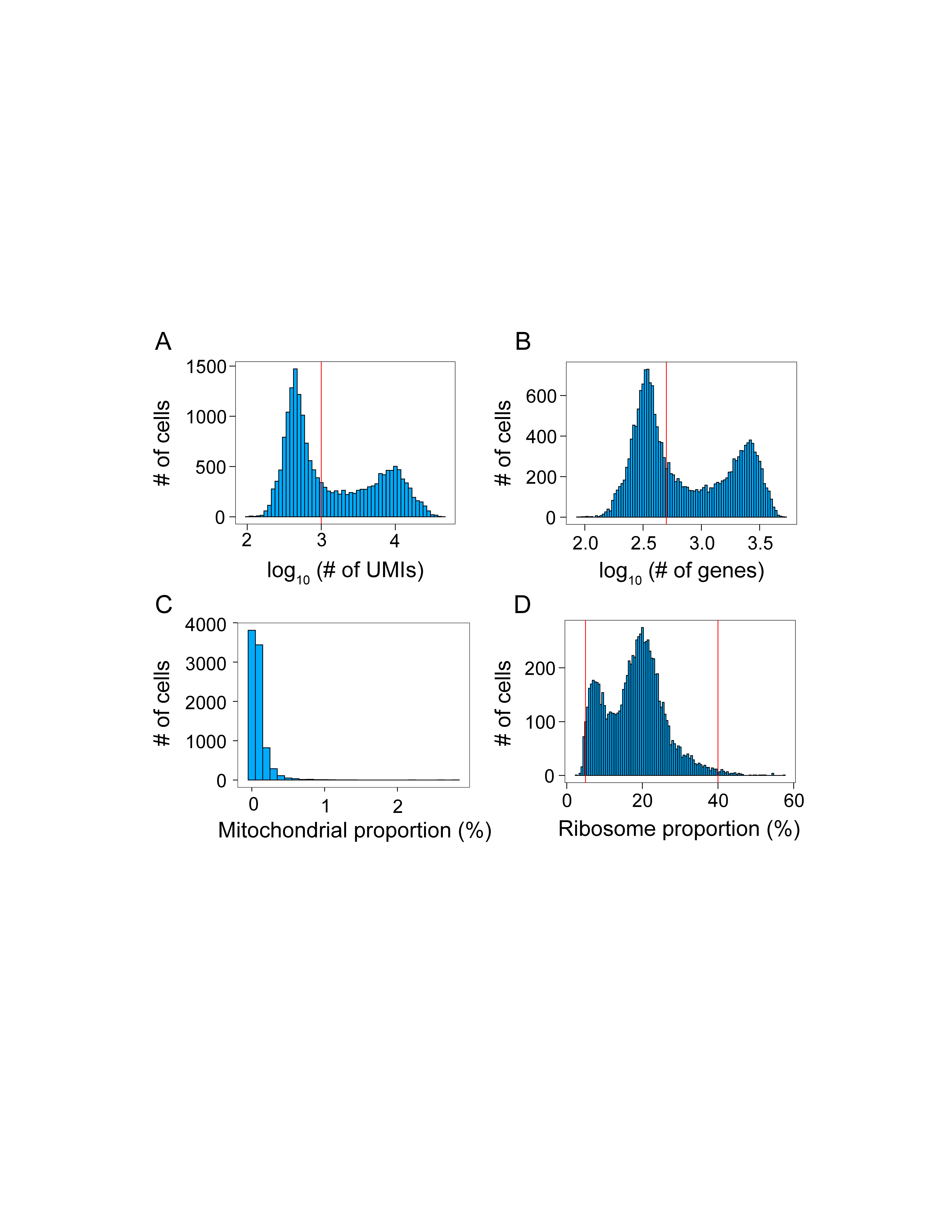

### Figure S2_ver5.png

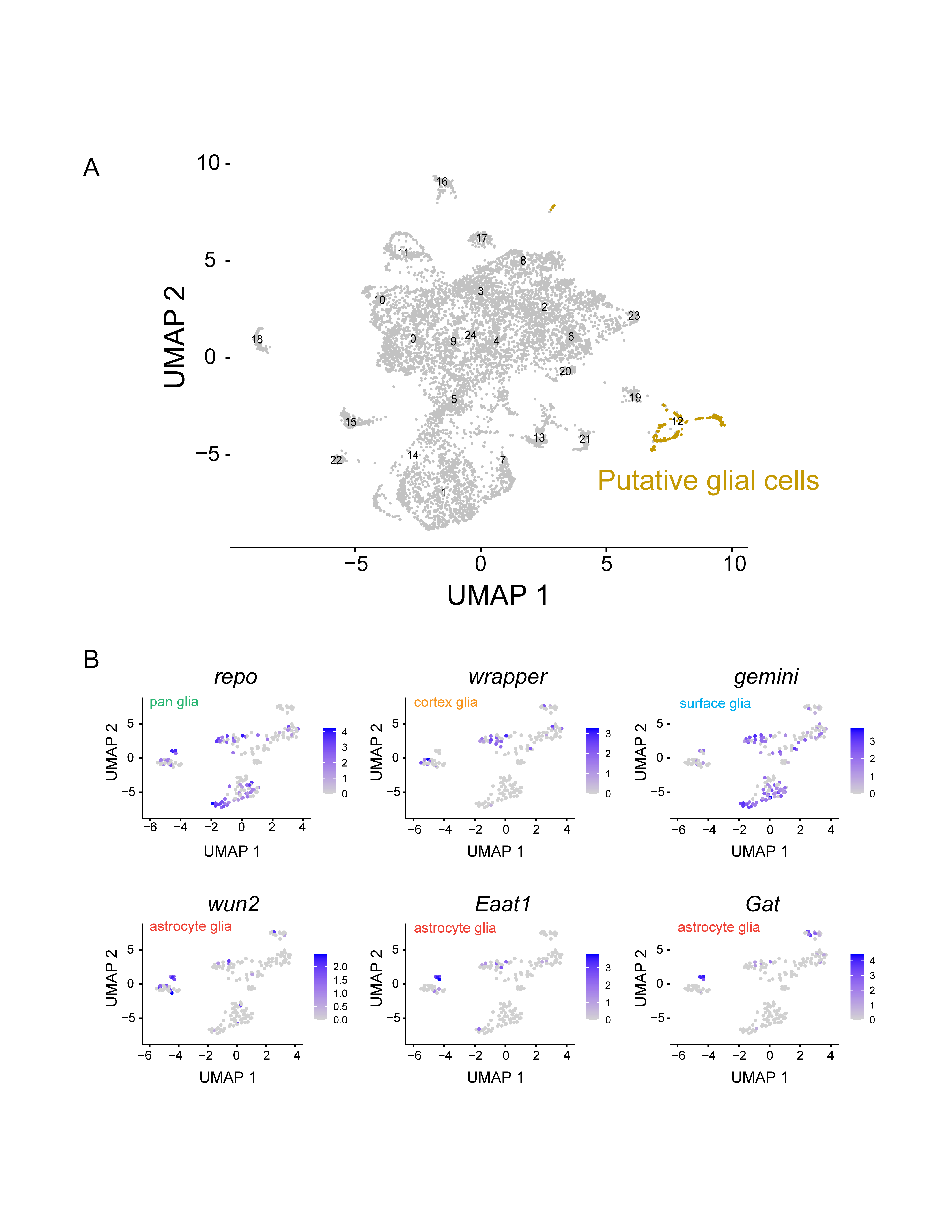

### Figure S3_ver5.png

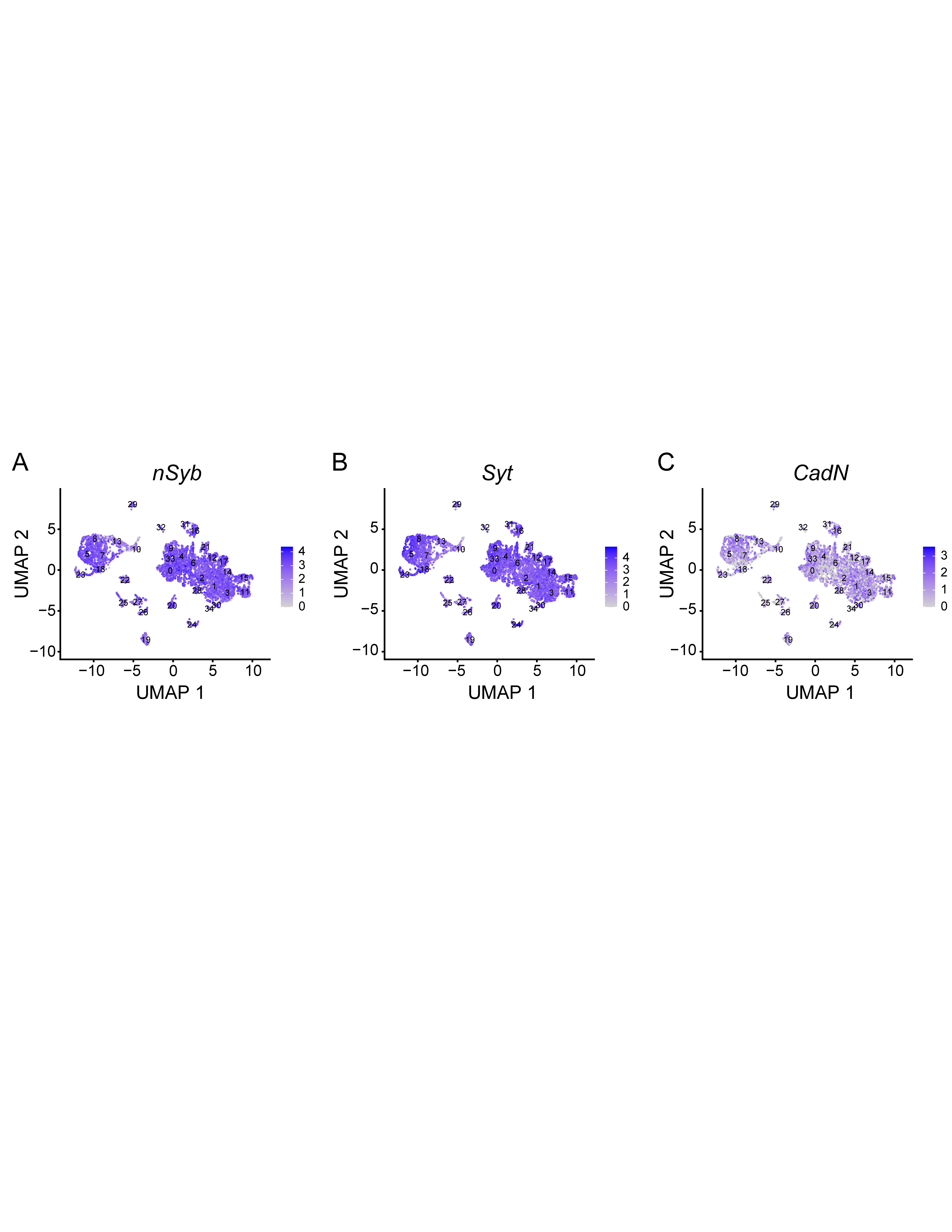

### Figure S4_ver5.png

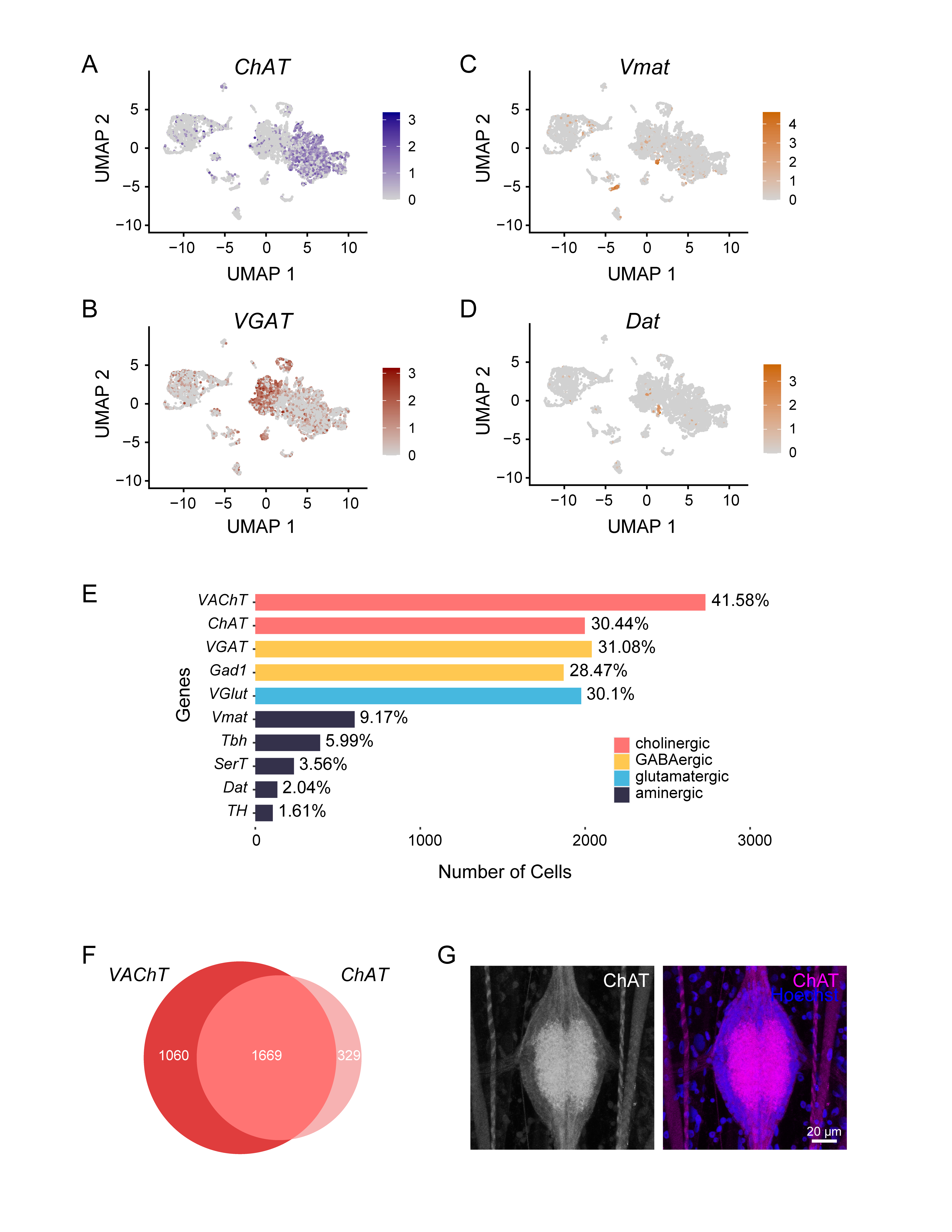

### Figure S5_ver5.png

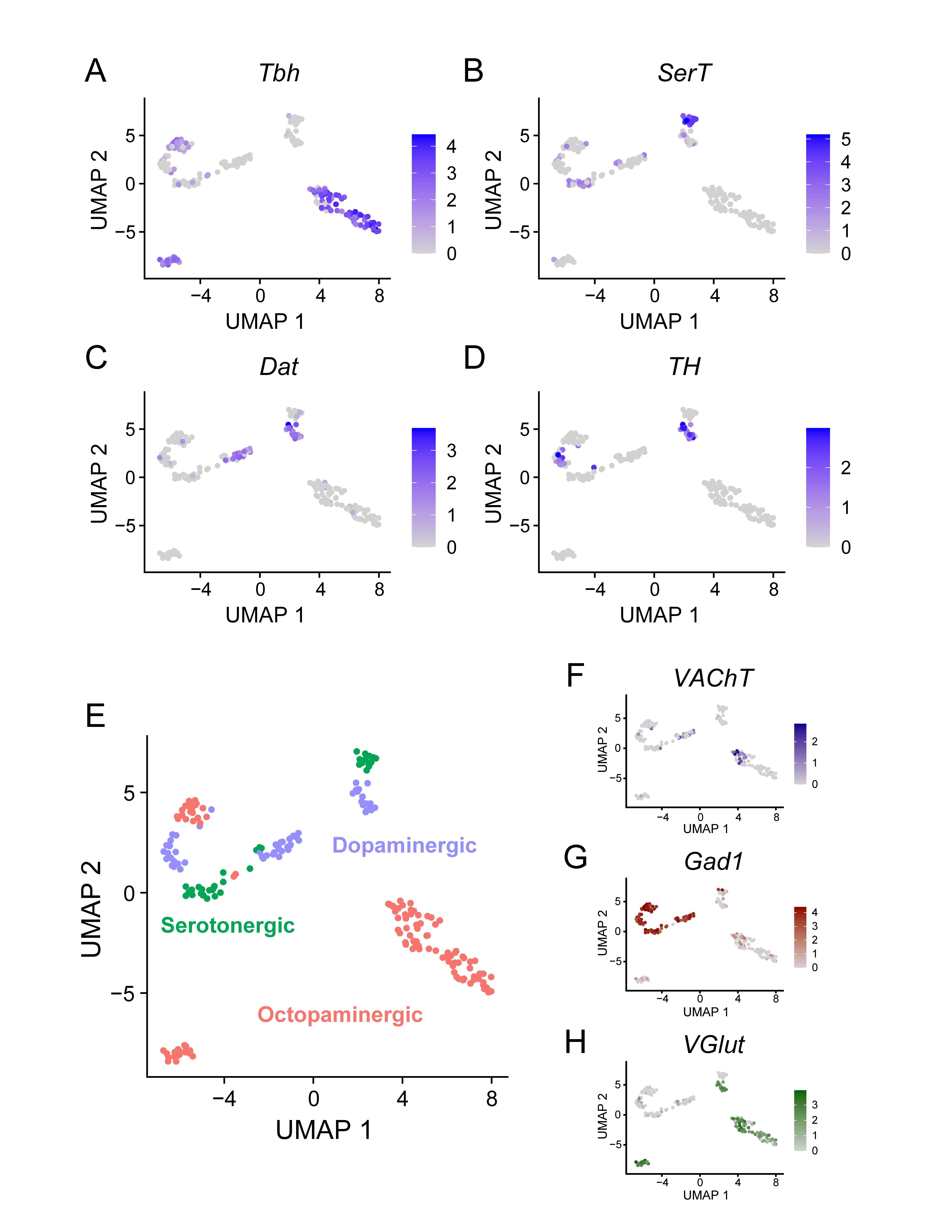

### Figure S6_ver5.png

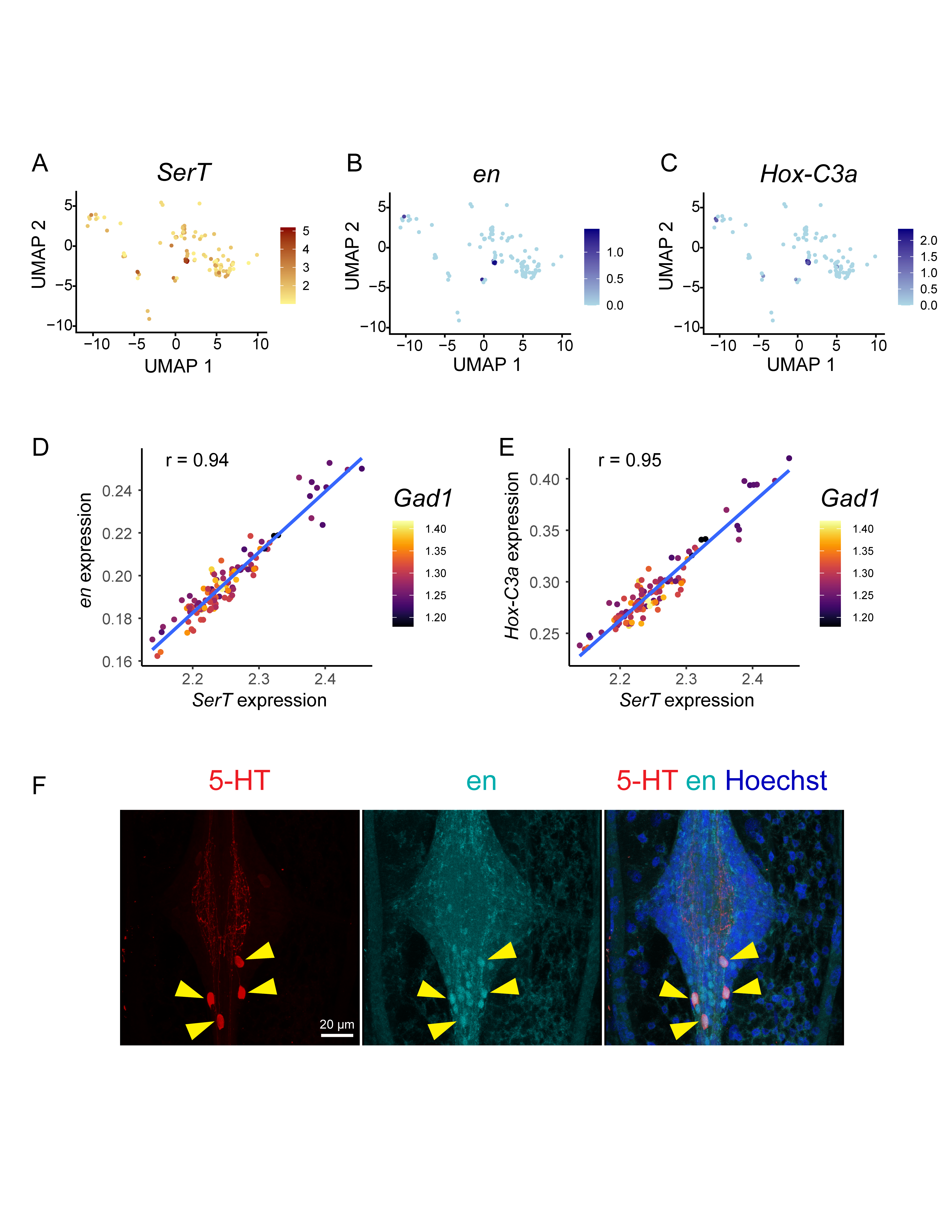

### Figure S7_ver5.png

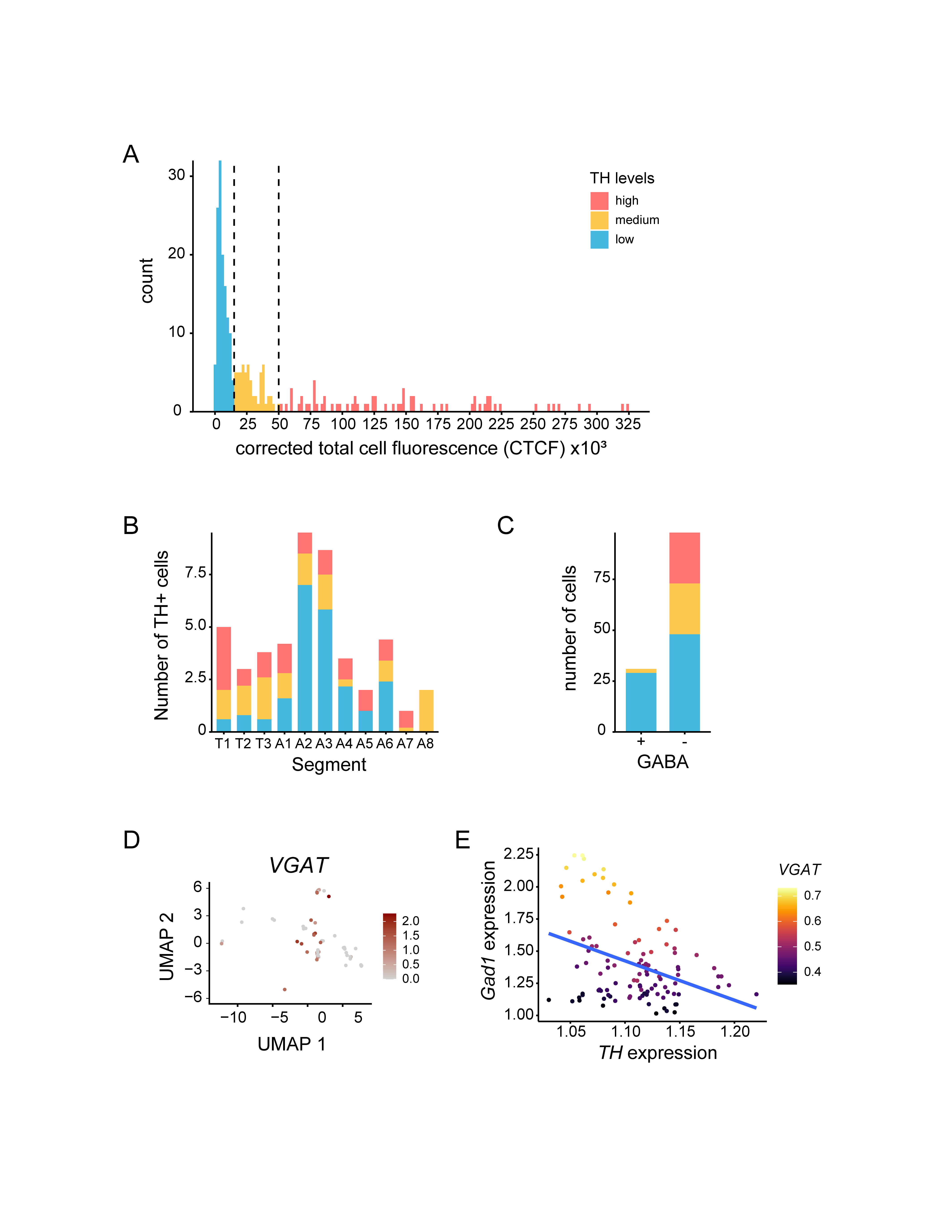

### Figure S8_ver5.png

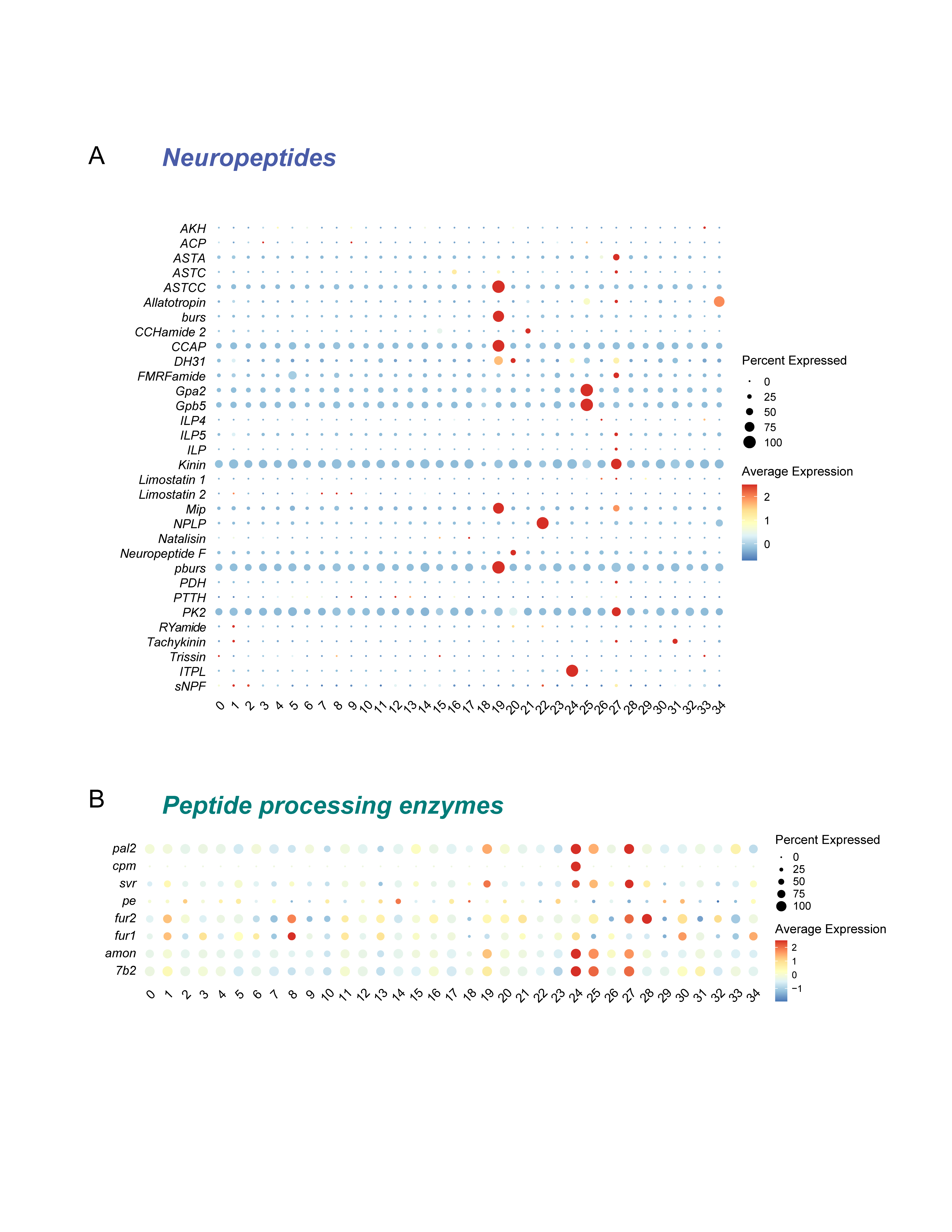

### Figure s9_ver5.png

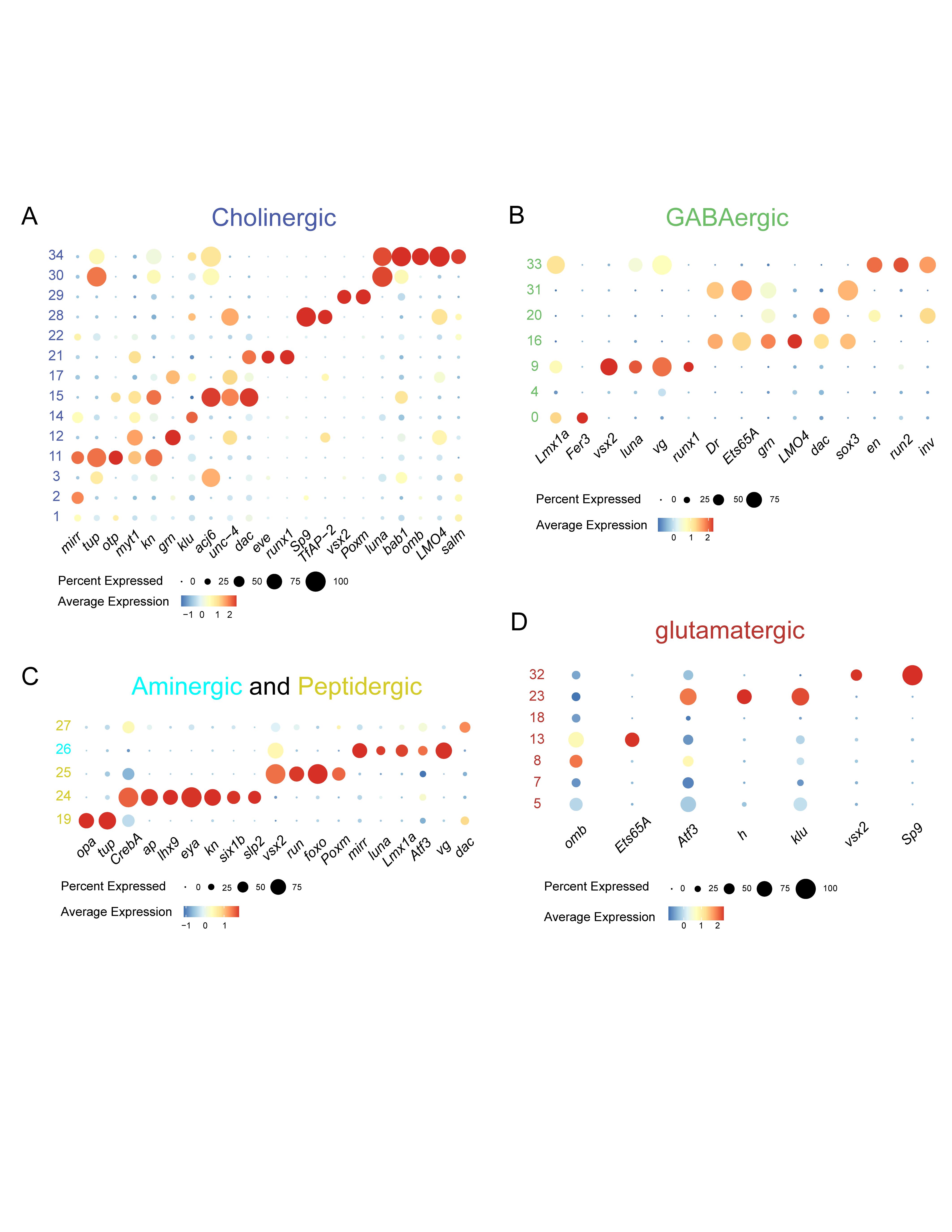

### Figure S10_ver5.png

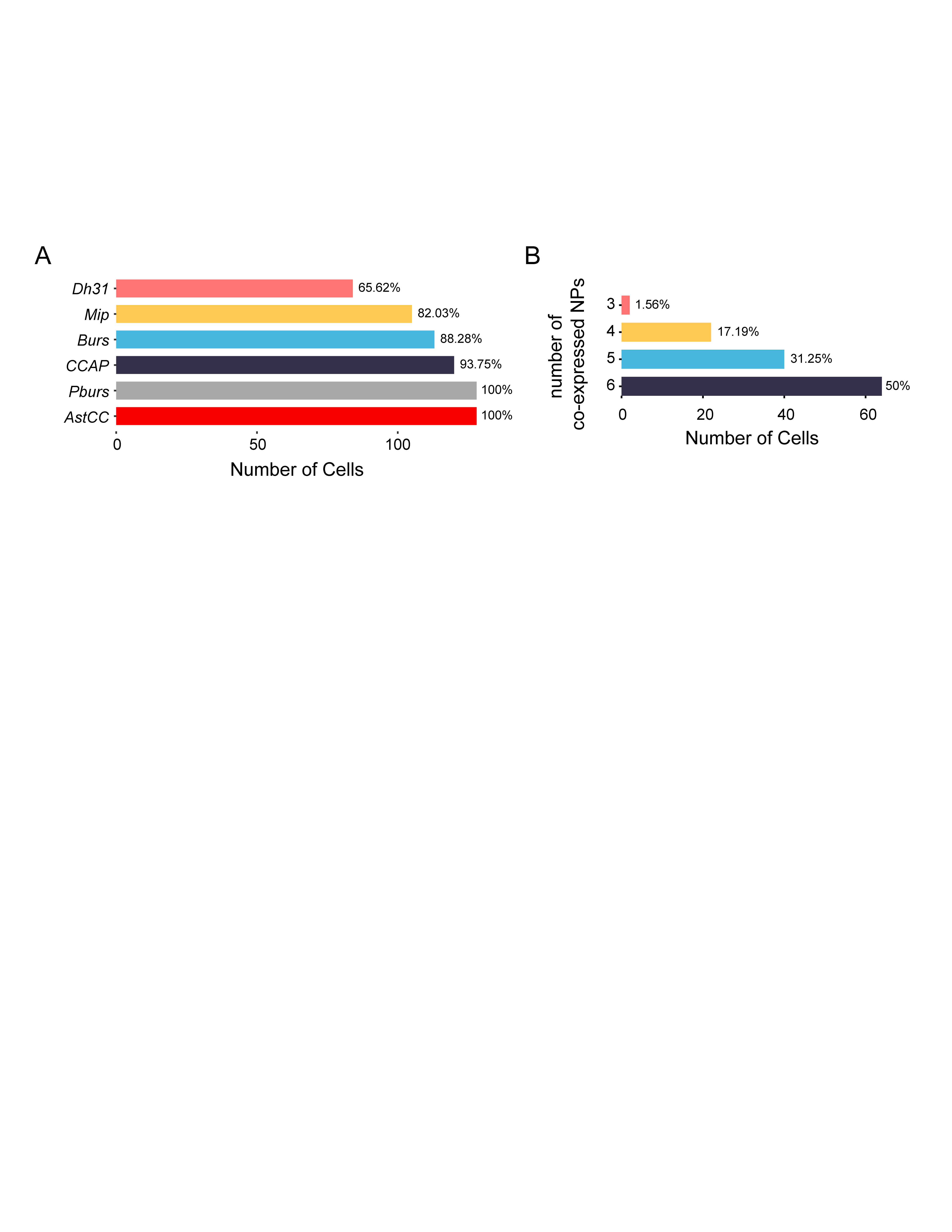

### Figure S11_ver5.png

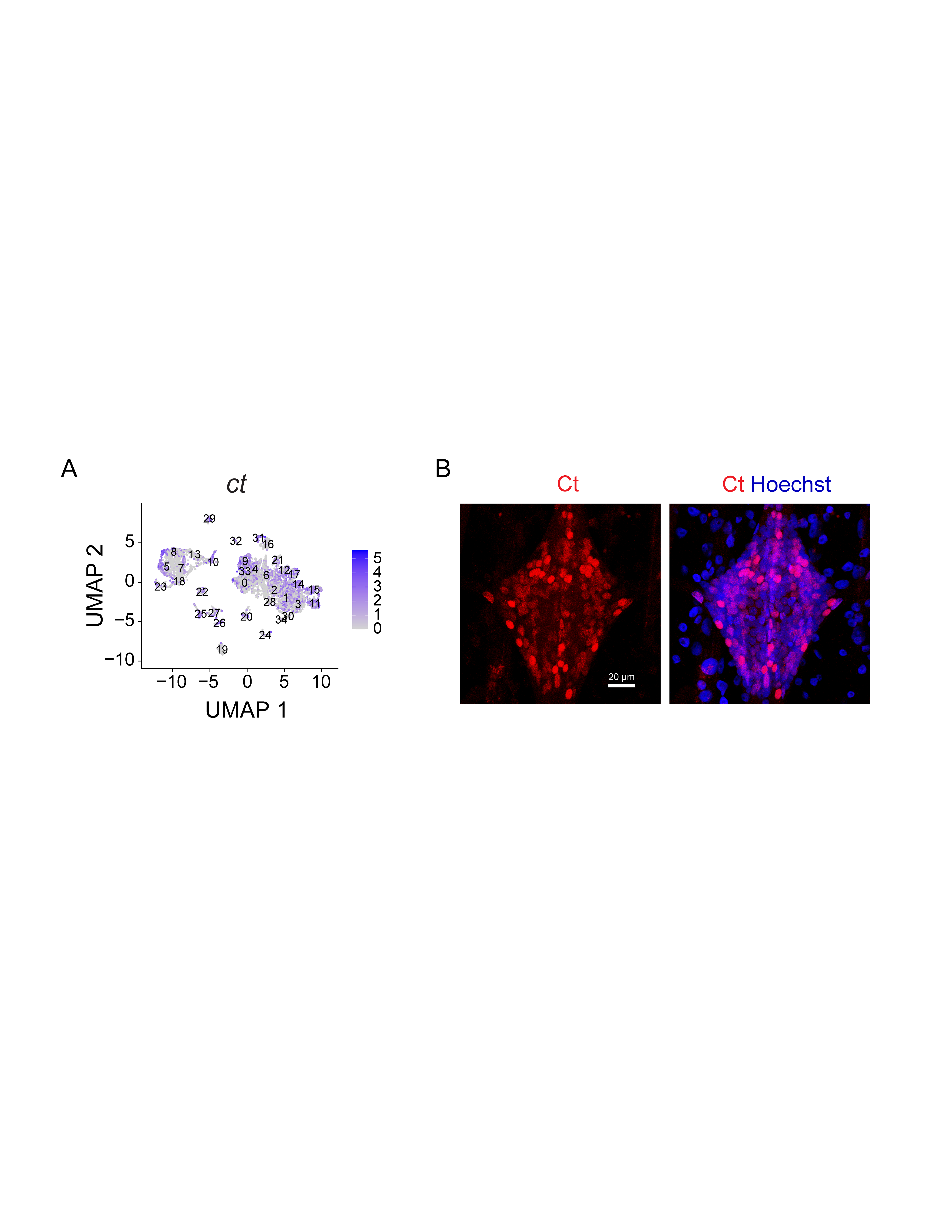

### Figure S12_ver5.png

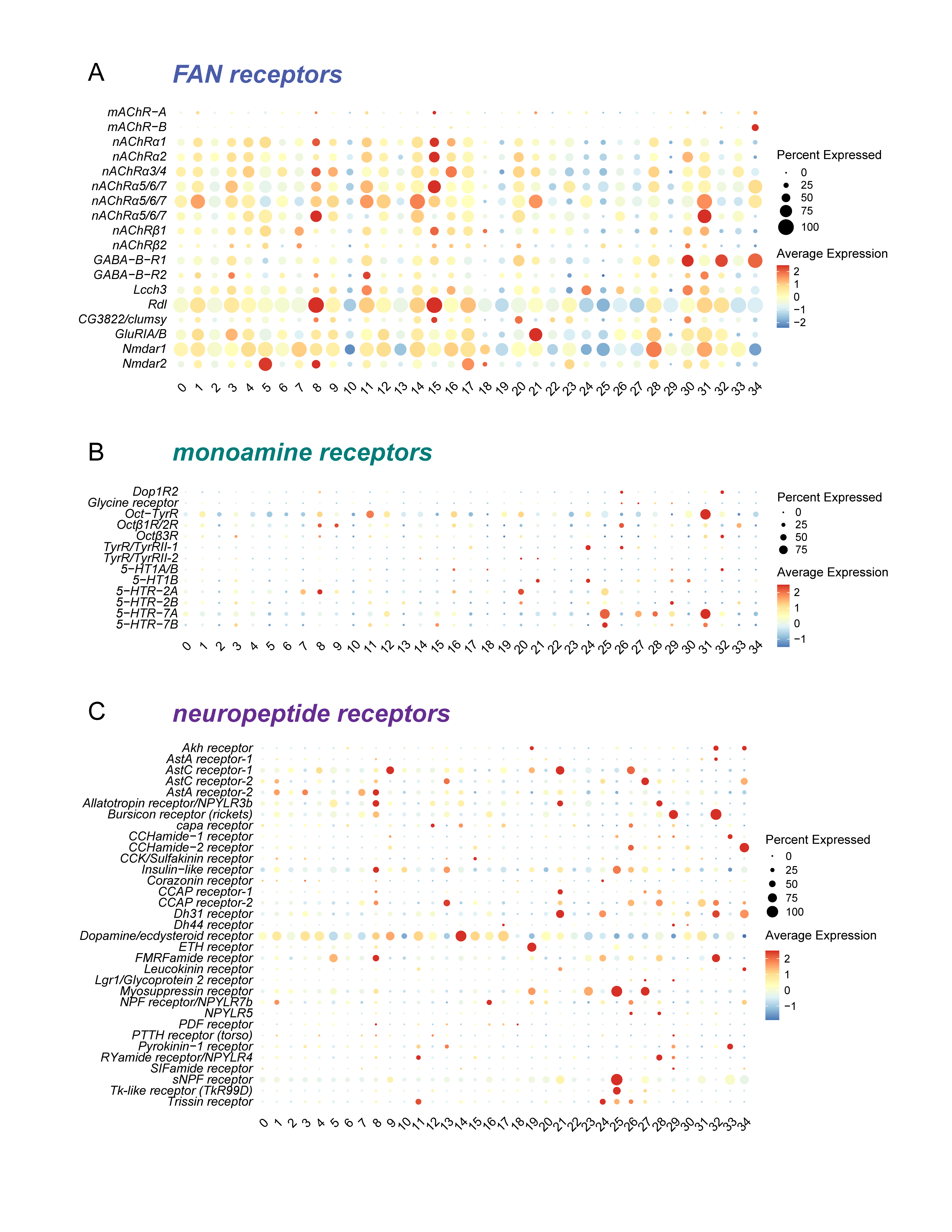

### Figure S13_ver5.png

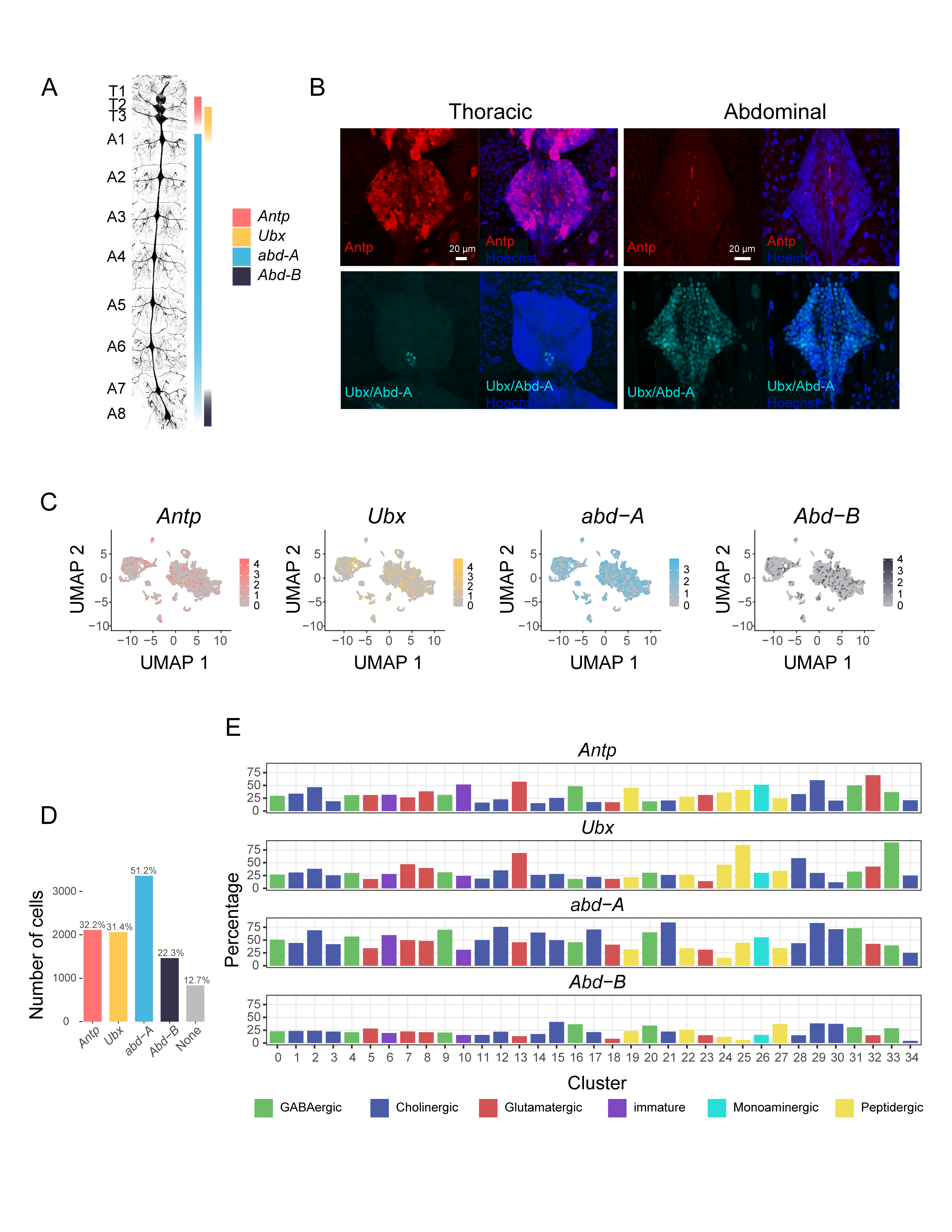

### Figure s14_ver5.png

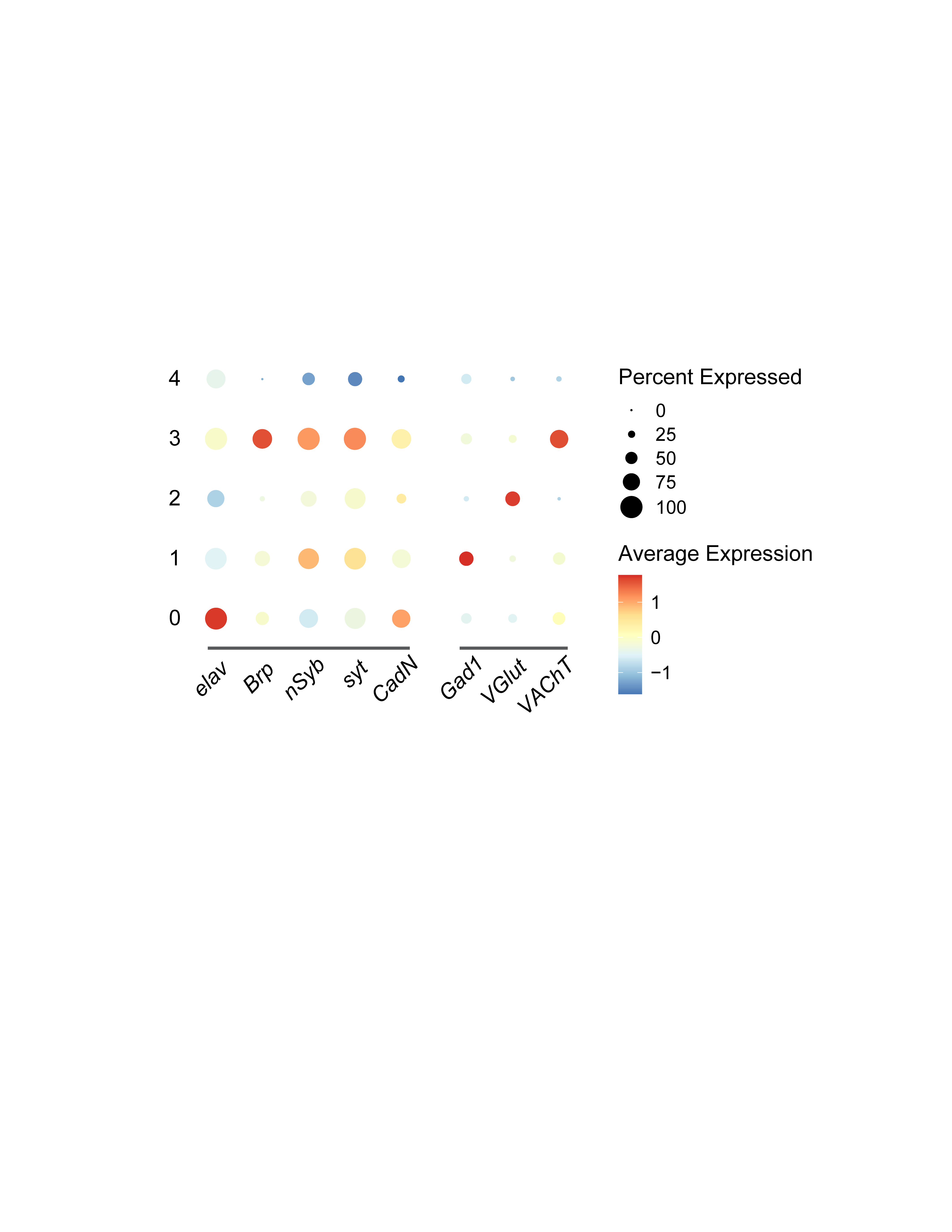

### Figure S15_ver4.png

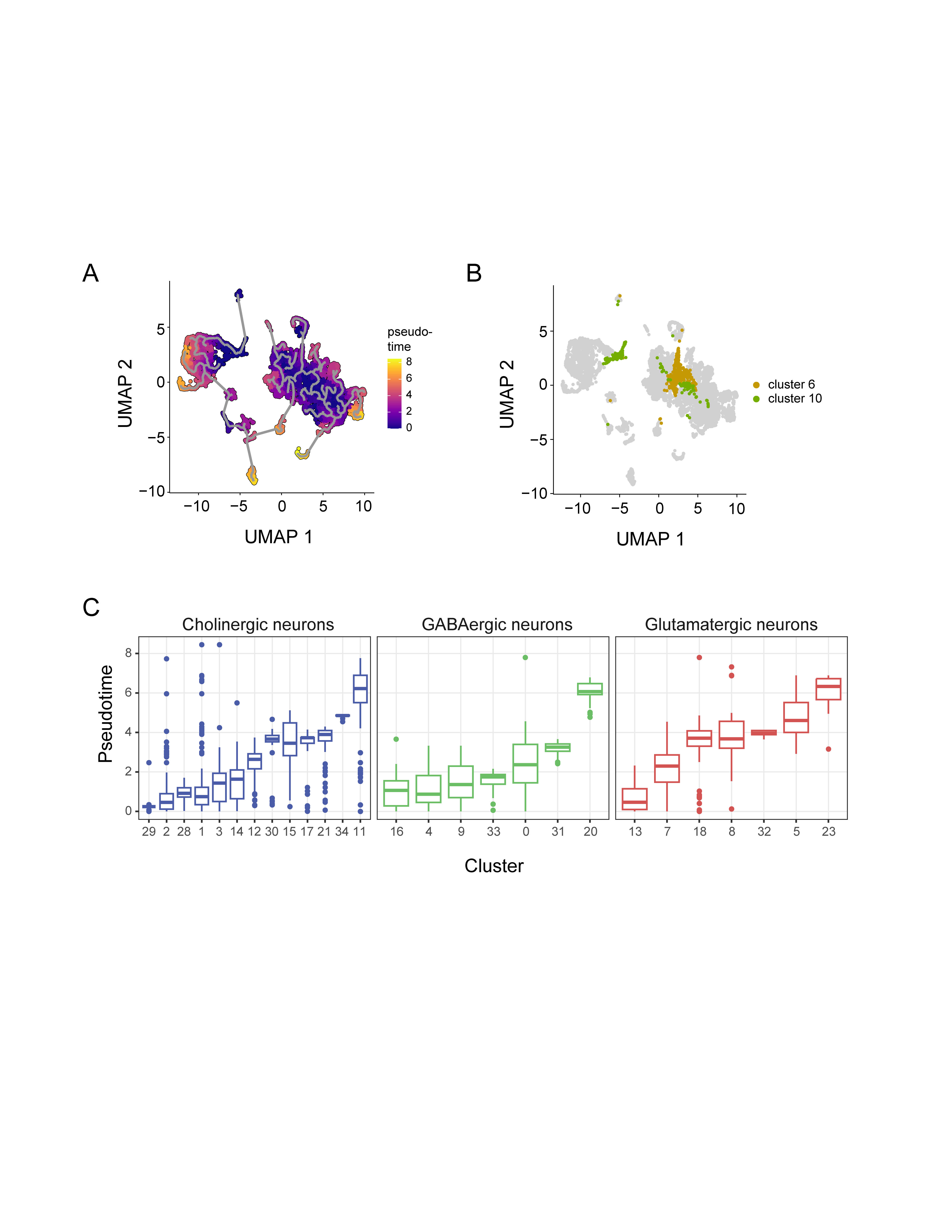

### Figure s16_ver5.png

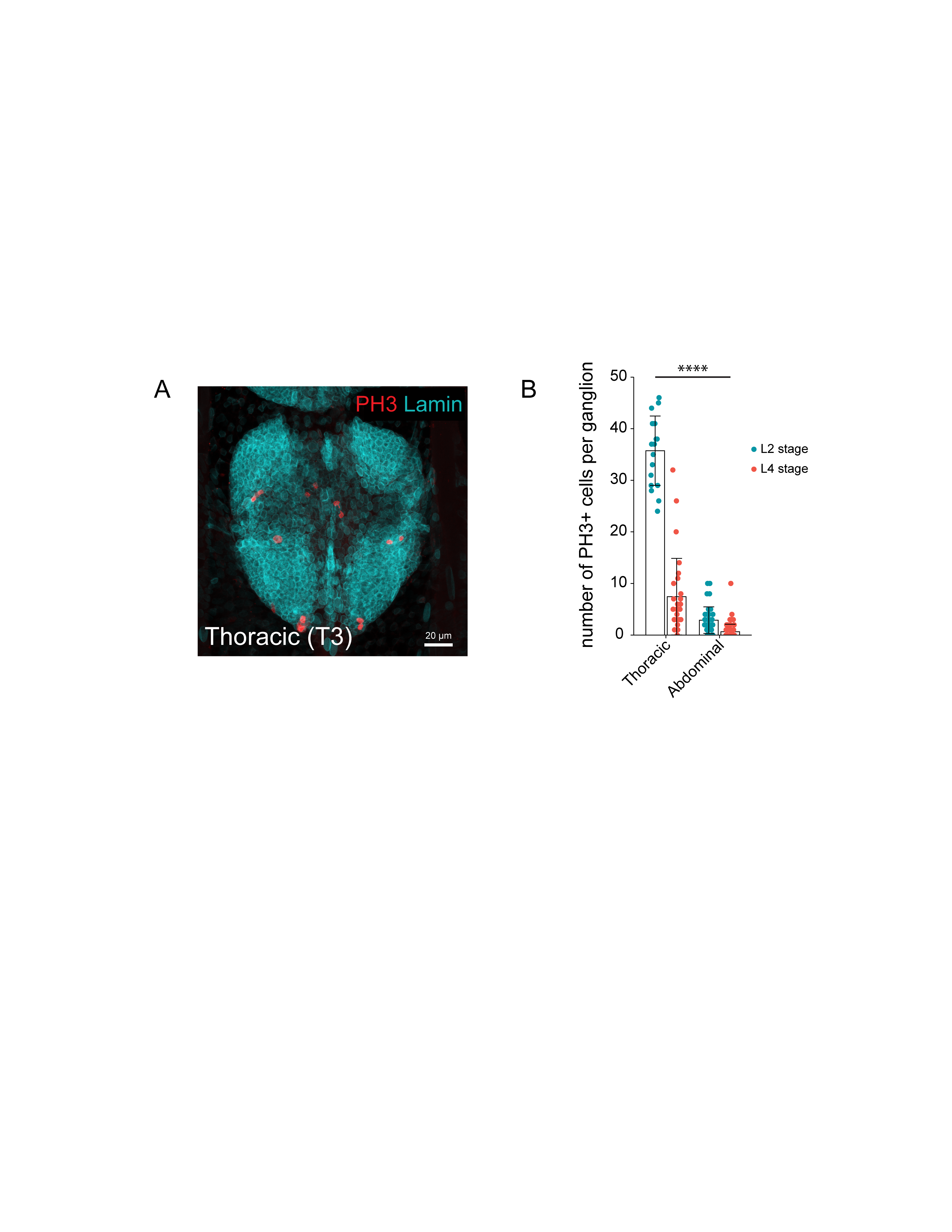

### FigureS17_ver5.png

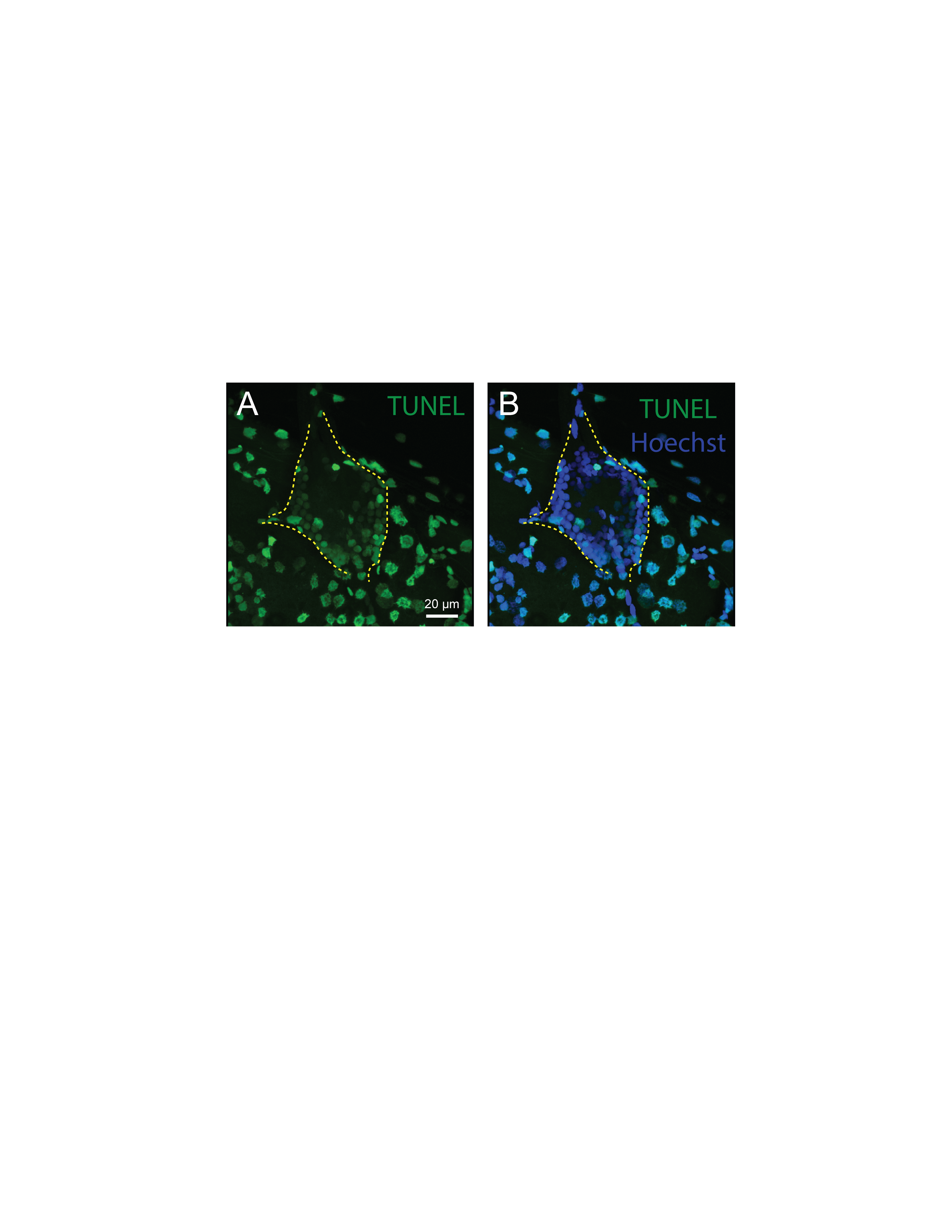
