## Supplemental Tables for "A cell atlas of the larval *Aedes aegypti* ventral nerve cord": YinAedesVNC_TableS3_NT_cluster.pdf

**Table S3. Proportion of cells in each cluster expressing neurotransmitter marker genes.**

| cluster | Cholinergic | GABAergic | Glutamatergic | Monoaminergic |
|---|---|---|---|---|
| 0 | 14.74% | 98.18% | 4.47% | 13.41% |
| 1 | 91.51% | 27.17% | 9.06% | 26.60% |
| 2 | 83.61% | 20.90% | 15.91% | 17.58% |
| 3 | 92.34% | 21.29% | 4.31% | 21.05% |
| 4 | 17.06% | 89.76% | 12.60% | 31.50% |
| 5 | 6.09% | 15.07% | 99.42% | 11.59% |
| **6** | 41.69% | 24.45% | 59.87% | 15.05% |
| 7 | 19.29% | 15.43% | 98.71% | 12.86% |
| 8 | 24.33% | 39.16% | 98.48% | 13.31% |
| 9 | 10.95% | 97.01% | 4.48% | 12.44% |
| **10** | 33.83% | 25.87% | 24.88% | 14.93% |
| 11 | 95.92% | 24.49% | 4.59% | 15.31% |
| 12 | 96.91% | 23.20% | 5.15% | 14.95% |
| 13 | 13.33% | 20.56% | 92.78% | 11.67% |
| 14 | 93.60% | 19.77% | 8.72% | 17.44% |
| 15 | 86.34% | 22.36% | 4.35% | 14.29% |
| 16 | 15.09% | 98.11% | 5.66% | 14.47% |
| 17 | 100.00% | 21.15% | 3.21% | 17.95% |
| 18 | 17.81% | 22.60% | 76.03% | 5.48% |
| **19** | 14.06% | 29.69% | 10.16% | 22.66% |
| 20 | 4.46% | 93.75% | 2.68% | 40.18% |
| 21 | 87.96% | 29.63% | 10.19% | 17.59% |
| **22** | 43.62% | 38.30% | 18.09% | 24.47% |
| 23 | 8.60% | 11.83% | 95.70% | 8.60% |
| **24** | 17.58% | 23.08% | 9.89% | 7.69% |
| **25** | 9.41% | 17.65% | 12.94% | 7.06% |
| 26 | 13.16% | 30.26% | 73.68% | 97.37% |
| **27** | 12.33% | 27.40% | 16.44% | 24.66% |
| 28 | 95.89% | 20.55% | 8.22% | 15.07% |
| 29 | 70.00% | 16.67% | 5.00% | 1.67% |
| 30 | 93.22% | 23.73% | 5.08% | 28.81% |
| 31 | 0.00% | 98.08% | 3.85% | 34.62% |
| 32 | 10.00% | 30.00% | 97.50% | 50.00% |
| 33 | 7.89% | 100.00% | 5.26% | 15.79% |
| 34 | 95.83% | 29.17% | 0.00% | 33.33% |
