## Supplemental Tables for "A cell atlas of the larval *Aedes aegypti* ventral nerve cord": YinAedesVNC_TableS4_NT_coExp.pdf

**Table S4. Proportion of cells in each cluster expressing marker genes for 0, 1, or multiple neurotransmitters.**

| cluster | 0 | 1 | 2 | 3 | 4 |
|---|---|---|---|---|---|
| 0 | 0.33% | 71.52% | 25.33% | 2.65% | 0.17% |
| 1 | 3.21% | 49.43% | 37.55% | 9.43% | 0.38% |
| 2 | 2.38% | 62.71% | 29.93% | 4.51% | 0.48% |
| 3 | 3.59% | 59.81% | 30.86% | 5.50% | 0.24% |
| 4 | NA | 55.64% | 38.32% | 5.51% | 0.52% |
| 5 | 0.58% | 70.72% | 24.64% | 4.06% | NA |
| 6 | 0.94% | 62.70% | 30.72% | 5.64% | NA |
| 7 | 0.64% | 61.09% | 29.58% | 8.68% | NA |
| 8 | 0.76% | 42.21% | 39.16% | 16.73% | 1.14% |
| 9 | 2.49% | 71.64% | 24.38% | 1.49% | NA |
| **10** | **25.37%** | 52.24% | 20.40% | 1.49% | 0.50% |
| 11 | 2.55% | 58.67% | 34.69% | 4.08% | NA |
| 12 | 2.58% | 58.25% | 35.57% | 3.61% | NA |
| 13 | 4.44% | 58.89% | 30.56% | 6.11% | NA |
| 14 | 1.74% | 62.79% | 29.65% | 5.81% | NA |
| 15 | 6.21% | 63.35% | 27.33% | 3.11% | NA |
| 16 | 1.89% | 67.30% | 26.42% | 4.40% | NA |
| 17 | NA | 64.74% | 28.85% | 5.77% | 0.64% |
| 18 | 13.01% | 55.48% | 28.08% | 3.42% | NA |
| **19** | **41.41%** | 44.53% | 10.16% | 3.91% | NA |
| 20 | 5.36% | 50.00% | 42.86% | 1.79% | NA |
| 21 | 5.56% | 51.85% | 36.11% | 4.63% | 1.85% |
| **22** | **19.15%** | 44.68% | 28.72% | 7.45% | NA |
| 23 | 2.15% | 74.19% | 20.43% | 3.23% | NA |
| **24** | **54.95%** | 32.97% | 10.99% | 1.10% | NA |
| **25** | **60.00%** | 32.94% | 7.06% | NA | NA |
| 26 | 1.32% | 15.79% | 55.26% | 22.37% | 5.26% |
| **27** | **43.84%** | 32.88% | 21.92% | 1.37% | NA |
| 28 | 4.11% | 64.38% | 20.55% | 9.59% | 1.37% |
| **29** | **21.67%** | 63.33% | 15.00% | NA | NA |
| 30 | 1.69% | 57.63% | 30.51% | 8.47% | 1.69% |
| 31 | 1.92% | 59.62% | 38.46% | NA | NA |
| 32 | 2.50% | 27.50% | 50.00% | 20.00% | NA |
| 33 | NA | 76.32% | 21.05% | NA | 2.63% |
| 34 | NA | 45.83% | 50.00% | 4.17% | NA |
